# Discovery of the β-Carboline Biosynthetic Pathway Enables *In Planta* Reconstruction of the Amazonian Brew Ayahuasca

**DOI:** 10.64898/2026.11.12.766139

**Authors:** Let Kho Hao, Shahar Cohen, Janka Höfer, Bekele Abebie Tuffa, Olga Khersonsky, Annu Kumari, Efrat Almekias-Siegl, Adi Faigenboim, Younghui Dong, Uwe Heinig, Sagit Meir, Ilana Rogachev, Itay Maoz, Sarel-Jacob Fleishman, Asaph Aharoni, Paula Berman

## Abstract

β-Carbolines are bioactive indole alkaloids with remarkable pharmacological properties and a defining role in ayahuasca, a traditional Amazonian shamanic brew, where their inhibition of monoamine oxidases enables the oral activity of the psychoactive *N,N*-dimethyltryptamine (DMT). Yet, despite their longstanding medicinal and cultural significance, the biosynthetic origin of β-carbolines in plants remains largely unresolved. Here, we uncover key steps underlying the biosynthesis of harmine, the major bioactive β-carboline alkaloid in *Peganum harmala*, by integrating spatial metabolite and transcriptome profiling with enzyme characterization and pathway reconstruction. We discovered four enzymes that enable *de novo* harmine biosynthesis in *Nicotiana benthamiana* and revealed how two closely related cytochrome P450s exhibit divergent regioselectivity, directing tryptamine towards either β-carboline or serotonin biosynthesis. Structural modeling and targeted mutagenesis pinpoint active-site residues that switch this regioselectivity, revealing a molecular basis for diversification of tryptamine metabolism. By combining distinct tryptamine pathways, we achieved *de novo* co-production of β-carbolines and DMT in *N. benthamiana* and *P. harmala* hairy roots without supplementing exogenous precursors. Together, these findings unlock β-carboline biosynthesis for future sustainable bioproduction with broad opportunities spanning medicine, crop protection, and biodiversity conservation.

## Introduction

β-Carboline alkaloids are a diverse class of tryptophan-derived specialized metabolites with a broad range of pharmacological activities, including antidepressant, anti-inflammatory, neuroprotective, antidiabetic and antitumor effects (de Osório et al., 2015; Zhang et al., 2020; Aaghaz et al., 2021). In plants, β-carbolines have been implicated in ecological interactions across trophic levels, including anti-herbivore and antimicrobial activities and phytotoxic effects on other plants, such as inhibition of root growth and disruption of cell division (Cavin and Rodriguez, 1988; Bouayad et al., 2012; Shao et al., 2013; Cui et al., 2020; Diehl et al., 2026). They are found across multiple plant lineages and occur in distantly related plant families, raising the possibility that distinct biosynthetic pathways may have evolved to produce structurally related alkaloids (Patel et al., 2012; Dai et al., 2018). Among β-carboline-producing plants, *Peganum harmala* (*P. harmala*, Nitrariaceae) is native to arid and semi-arid regions of the Mediterranean, Middle East, and Central Asia (Sharifi-Rad et al., 2021). It has a long history of medicinal and ritual use, which has attracted considerable interest for its diverse pharmacological and psychoactive properties (Moloudizargari et al., 2013).

Harmine and harmaline were among the earliest characterized β-carboline (harmala) alkaloids, having been isolated from *P. harmala* seeds in the nineteenth century (Fritzsche, 1848). Subsequent phytochemical studies identified additional β-carbolines in *P. harmala*, with harmalol, tetrahydroharmol (THH-OH), and tetrahydroharmine (THH) detected predominantly in seeds, and harmol particularly abundant in roots (Herraiz et al., 2010; Fang et al., 2019). These alkaloids are also central to ayahuasca, a traditional Amazonian psychoactive brew, in which β-carboline-rich *Banisteriopsis caapi* (*B. caapi*, Malpighiaceae) is combined with *N*,*N*-dimethyltryptamine (DMT)-producing *Psychotria viridis* (Santos et al., 2020). In this preparation, β-carboline-mediated inhibition of monoamine oxidase A (MAO-A) enables the oral bioavailability of DMT. *Peganum harmala* has likewise been used as a β-carboline source in ayahuasca analogue preparations (Kaasik et al., 2021). Growing interest in the therapeutic potential of psychedelics has stimulated clinical investigations of ayahuasca, particularly for treatment-resistant depression and other psychiatric disorders (Palhano-Fontes et al., 2019; Dos Santos and Hallak, 2024). Despite this growing interest, the biosynthetic pathways of β-carbolines in plants remain poorly characterized, even after decades of biochemical investigations.

Early isotope-labeling studies investigating harmine **1** biosynthesis in *P. harmala* identified L-tryptophan (Trp **2**) and tryptamine (TAM **3**) as precursors and demonstrated the incorporation of pyruvate **4** into its β-carboline skeleton (Stolle and Gröger, 1968; **Supplemental Figure S1A**). Subsequent feeding experiments in *P. harmala* callus cultures demonstrated the oxidation of harmaline **5** to harmine **1**, but failed to detect incorporation of 6-hydroxytryptamine (6-HT **6**), a proposed precursor of harmalol **7**, into β-carboline alkaloids (Nettleship and Slaytor, 1974). Nevertheless, the authors proposed a biosynthetic route involving 6-HT **6**, harmalol **7**, and harmaline **5** as successive intermediates leading to harmine **1**, with a possible alternative route through harmol **8** and its methylation to harmine **1**. They also proposed the biosynthesis of serotonin (5-hydroxytryptamine, 5-HT **9**) from Trp **2** *via* TAM **3** to explain its accumulation in *P. harmala* (Nettleship and Slaytor, 1974; **Supplemental Figure S1B**). More recently, Jamieson et al. (2021) proposed that the β-carboline scaffold results from Pictet-Spengler condensation of TAM **3** with pyruvate **4**, yielding 1,2,3,4-tetrahydro-β-carboline-1-carboxylic acid (THβC-1-COOH **10**), followed by oxidative decarboxylation to 1-methyl-β-carboline **11 (Supplemental Figure S1C**). Similar Pictet-Spengler reactions could also proceed with 6-HT **6** instead of TAM **3**, or acetaldehyde **12** could replace pyruvate **4** as the carbonyl donor, bypassing the carboxylated tetrahydro-β-carboline intermediate (Callaway et al., 1994; **Supplemental Figure S1D**).

In this study, we elucidated key steps in the biosynthesis of harmine **1** and related β-carbolines from Trp **2** in *P. harmala*, addressing unresolved steps from the earlier precursor-feeding studies. We identified two closely related cytochrome P450s (CYPs) that direct TAM **3** hydroxylation toward 5-HT **9** or 6-HT **6**, establishing 6-HT **6** as the entry point to β-carboline biosynthesis. Structural modeling and targeted mutagenesis further identified two amino acid residues whose reciprocal substitution switched the regioselectivities of these CYPs. We next demonstrated that acetaldehyde **12** serves as a carbonyl donor in a nonenzymatic Pictet-Spengler-type reaction with 6-HT **6** and identified an *O*-methyltransferase (OMT) active on multiple β-carboline intermediates. Consequently, we reconstructed β-carboline biosynthesis in *Nicotiana benthamiana* (*N. benthamiana*), with some steps occurring spontaneously or potentially mediated by endogenous host enzymes, and achieved simultaneous production of β-carbolines and DMT **13** through transient expression. Finally, we established a *P. harmala* hairy-root system engineered to express DMT **13** biosynthetic genes, enabling its accumulation alongside endogenous β-carbolines and recapitulating the principal alkaloid combination found in ayahuasca. This system offers future opportunities to decipher the β-carboline pathway compartmentalization and metabolite transport, while providing a platform for engineering the production of diverse TAM-derived metabolites.

## Results

### Harmine accumulates predominantly in the epidermis of young *P. harmala* roots

We first quantified harmine **1** across tissues of *P. harmala* and *B. caapi* by high-resolution liquid chromatography-tandem mass spectrometry (LC-MS/MS). Harmine **1** was detected in all analyzed tissues of both species, with the highest levels found in young roots of *P. harmala*, reaching 2.06 ± 0.25% of fresh weight (**Figure 1A** and **1B**, and **Supplemental Table S1** and **S2**). Using matrix-assisted laser desorption/ionization-mass spectrometry imaging (MALDI-MSI), we further found that harmine **1** was enriched in the pith and epidermis of *P. harmala* roots, whereas Trp **2** was observed predominantly in the cortex and phloem, suggesting that the conversion of Trp **2** to β-carbolines within roots may be spatially restricted by the differential distribution of biosynthetic enzymes (**Figure 1C**). The yellow coloration observed in the optical images coincided with the regions of harmine **1** accumulation and may reflect the optical properties of its conjugated π-electron system. In young and mature stems of *B. caapi*, harmine **1** was enriched toward the outer bark, with higher accumulation observed in mature stems (**Supplemental Figure S2**).

**Figure 1.**
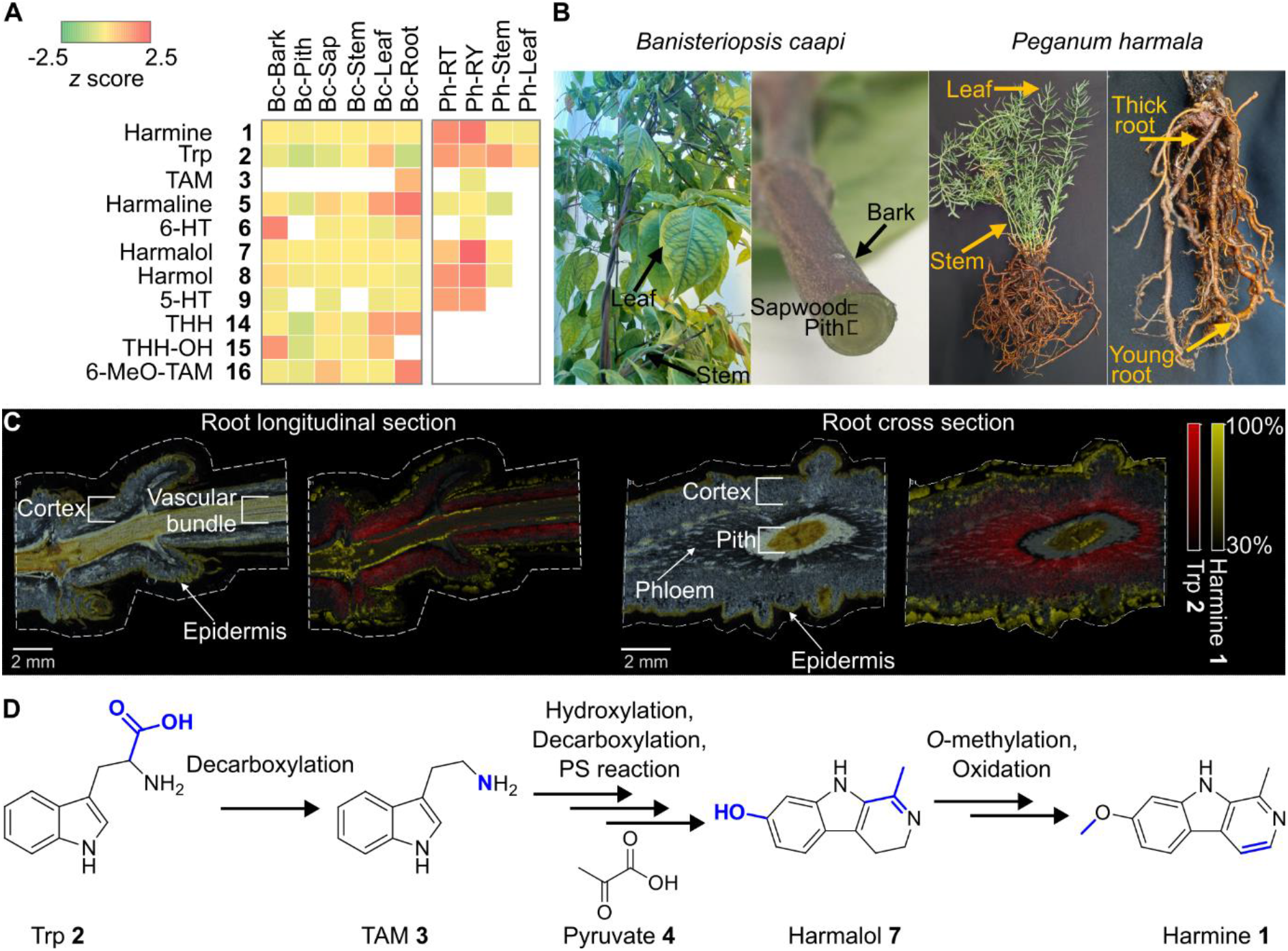
Tissue-specific distribution of β-carbolines. **(A)** Liquid chromatography-mass spectrometry (LC-MS) relative abundance of β-carbolines across *Banisteriopsis caapi* (*B. caapi*) and *Peganum harmala* (*P. harmala*) tissues (non-detected metabolites are shown in white; *z*-score; mean, *n* = 3 biological replicates). Metabolites were identified by LC-MS using analytical standards based on accurate mass, retention time, and MS/MS fragmentation patterns (**Supplemental Figure S3**). Tetrahydroharmol (THH-OH **15**) and 6-methoxy-tryptamine (6-MeO-TAM **16**) were tentatively identified based on accurate mass and comparison of their retention times and MS/MS fragmentation patterns with those of the structurally related compounds tetrahydroharmine (THH **14**) and 5-methoxy-tryptamine (**Supplemental Figure S4**). **(B)** Representative photographs of *B. caapi* and *P. harmala* plants and tissues used for metabolite profiling. **(C)** Matrix-assisted laser desorption/ionization-mass spectrometry imaging (MALDI-MSI) of *P. harmala* root sections (*n* = 2 biological replicates) showing the spatial distribution of harmine **1** ([M+H]^+^, m/z 213.102 ± 0.01 Da) and tryptophan (Trp **2**, [M+H]^+^, m/z 205.097 ± 0.01 Da). **(D)** Simplified predicted biosynthetic pathway of β-carbolines in *P. harmala.* Several arrows represent multiple reaction steps. The continuation of Figure 1 appears on the next page. TAM **3**, tryptamine; 6-HT **6**, 6-hydroxy-tryptamine; 5-HT **9**, 5-hydroxy-tryptamine; THβC-1-COOH **10**, 1,2,3,4-tetrahydro-β-carboline-1-carboxylic acid; THH **14**, tetrahydroharmine; THH-OH **15**, tetrahydroharmol; 6-MeO-TAM **16**, 6-methoxy-tryptamine. The legend of Figure 2 appears on the next page.

We further performed targeted profiling of β-carbolines and related metabolites in both species, focusing on previously identified metabolites and proposed intermediates of the β-carboline biosynthetic pathway (Aslam et al., 2014; Santos et al., 2020; Fahmy et al., 2021) (**Figure 1A**, **Supplemental Figure S3**, and **Supplemental Table S1**). Both species accumulated the β-carbolines harmalol **7**, harmol **8**, harmaline **5**, and harmine **1**. In *P. harmala*, these metabolites accumulated predominantly in roots, whereas in *B. caapi*, most showed a relatively uniform distribution across tissues, with higher levels of harmaline **5** and THH **14** detected in roots and leaves (**Figure 1A**). Interestingly, THH **14**, THH-OH **15**, and 6-methoxy-tryptamine (6-MeO-TAM **16**) were detected solely in *B. caapi* tissues (**Supplemental Figure S4** and **Supplemental Table S1**). 6-HT **6** was detected in all analyzed tissues of *B. caapi* except the stem pith, with highest accumulation in the outer stem area (bark), whereas in *P. harmala*, it was detected only in roots and at substantially lower levels (**Figure 1A**). In contrast, *P. harmala* exhibited high levels of 5-HT **9** in the roots, consistent with previous reports (McKenzie et al., 1975; Sasse et al., 1982; Berlin et al., 1993) (**Figure 1A**). Together, these results revealed both shared and species-specific features of β-carboline metabolism, potentially reflecting evolutionary divergence in the underlying biosynthetic pathways and their tissue-specific regulation. In this context, β-carboline-producing plants, including *P. harmala* and *B. caapi,* are placed in evolutionarily distant lineages, highlighting the broad phylogenetic distribution of these alkaloids and raising the possibility of convergent evolution of the β-carboline pathway (**Supplemental Figure S5** and **Supplemental Table S3**). Given the substantially higher harmine **1** accumulation and pronounced tissue-specific enrichment in *P. harmala*, we focused subsequent pathway-discovery efforts on this species. Based on our chemical profiling and previously proposed biosynthetic routes, we predicted a biosynthetic pathway for harmine **1** in *P. harmala*, in which Trp **2** is decarboxylated to TAM **3**, followed by hydroxylation, β-carboline scaffold formation, *O*-methylation, and oxidation steps (**Figure 1D** and **Supplemental Figure S1D**).

### Transcriptome-guided identification of key β-carboline biosynthetic enzymes

To identify β-carboline biosynthetic enzymes, we performed RNA sequencing of *P. harmala* leaves, stems, mature and young roots, matching the tissues used for metabolite profiling (**Figure 1A**). We generated *de novo* assemblies of transcriptomes using complementary TruSeq and Iso-Seq sequencing approaches to obtain comprehensive transcriptomic data in the absence of an annotated reference genome (**Supplemental Table S4**). To identify transcripts with tissue-specific expression patterns corresponding to the metabolite profiles, we performed hierarchical clustering of transcript expression profiles across the analyzed tissues. The resulting clusters revealed distinct tissue-specific expression patterns, including groups enriched in root tissues that corresponded to the preferential accumulation of β-carbolines (**Figure 2A**). Based on the predicted pathway (**Figure 1D** and **Supplemental Figure S1**), we focused on six candidate enzyme families: tryptophan decarboxylases (TDCs), CYP monooxygenases, potential Pictet-Spenglerases (PSs), pyruvate decarboxylases (PDCs), *O*-methyltransferases (OMTs), and flavin-dependent oxidoreductases. Candidate enzymes were prioritized according to their relative expression patterns across *P. harmala* tissues, together with predicted protein size, the presence of complete open reading frames (ORFs) and conserved Pfam domains, and sequence identity to previously characterized plant enzymes with similar activities.

**Figure 2.**
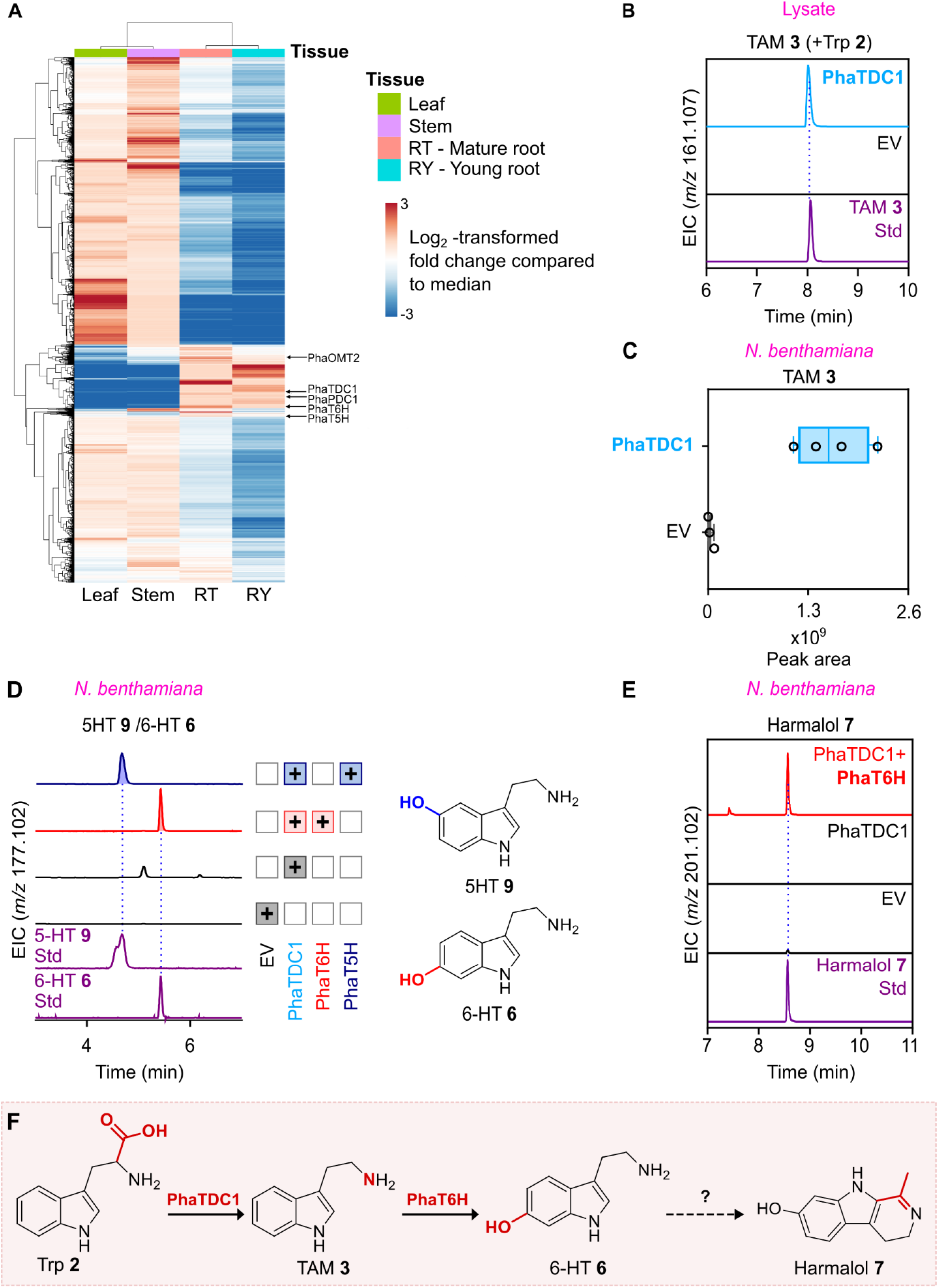
Functional characterization of tryptamine biosynthesis and hydroxylation in *Peganum harmala*. **(A)** Hierarchical clustering analysis was performed on trimmed mean of M-values (TMM)-normalized TruSeq transcripts using expression values converted to counts per million (CPM), with Euclidean distance and complete linkage. Transcripts with a CPM expression value greater than 25 in at least one of 12 tissue samples were retained (10,905 out of 307,954 total transcripts). Transcripts encoding enzymes characterized in this study are labeled with their names to illustrate their co-expression patterns. **(B)** Extracted ion chromatograms (EICs) of tryptamine (TAM **3**) following *in vitro* enzyme assays using *Escherichia coli* (*E. coli*) lysates expressing *PhaTDC1* with PLP and tryptophan (Trp **2**) as substrates (*n* = 3). **(C)** Ion abundance peak areas of TAM **3** following transient expression of *PhaTDC1* in *Nicotiana benthamiana* (*N. benthamiana*) leaves (median ± SD; *n* = 4 biological replicates). **(D)** EICs of 5-hydroxy-tryptamine (5-HT **9**) and 6-hydroxy-tryptamine (6-HT **6**) following transient co-expression of *PhaTDC1* with *PhaT6H* or *PhaT5H* in *N. benthamiana* leaves (*n* = 4). Plus signs (+) indicate biosynthetic genes expressed in each experiment, with boxes colored according to the last enzyme added in a set of coexpressed genes. Corresponding chemical structures are included for reference. **(E)** EICs of harmalol **7** following transient co-expression of *PhaTDC1* with *PhaT6H* (*n* = 4 biological replicates). **(F)** Schematic representation of 6-HT **6** biosynthesis and its conversion to harmalol **7** in *N. benthamiana* through a spontaneous or host enzyme-mediated reaction, as indicated by a question mark. All the metabolites were identified by LC-MS/MS using analytical standards based on accurate mass, retention time, and MS/MS fragmentation patterns. Chromatograms were normalized to the highest values. EV, empty vector; Std, standard.

### Regioselective tryptamine hydroxylation directs β-carboline biosynthesis

We identified a pyridoxal 5’-phosphate (PLP)-dependent decarboxylase candidate (*TDC1*) that was highly expressed in young roots and lowly expressed in aerial tissues (**Supplemental Table S5**). *In vitro* assays using *Escherichia coli* (*E. coli*) lysates expressing recombinant TDC1, supplemented with PLP and Trp **2**, demonstrated that TDC1 (PhaTDC1) catalyzed the decarboxylation of Trp **2** to TAM **3** (**Figure 2B**). We further confirmed PhaTDC1 *in vitro* activity using recombinant protein purified by nickel-affinity chromatography **(Supplemental Figure S6A-S6C**). Transient expression of *PhaTDC1* in *N. benthamiana* by agro-infiltration resulted in substantial accumulation of TAM **3** from endogenous Trp **2**, without exogenous substrate supplementation (**Figure 2C**). Using *PhaTDC1* as a reference gene, we further performed co-expression analysis to identify candidates for the subsequent biosynthetic steps. Applying a Pearson correlation coefficient (*r*) > 0.8 and false discovery rate (FDR)-adjusted *p* < 0.05 as thresholds, we identified 1,013 Iso-Seq and 176 TruSeq transcripts co-expressed with *PhaTDC1*, including 15 CYPs, 3 PDCs, 3 PSs, 15 OMTs, and 10 oxidases (**Supplemental Figure S6D** and **Supplemental Table S5**).

Several plant CYPs have been shown to catalyze the hydroxylation of TAM **3** to 5-HT **9** (Fujiwara et al., 2010; Negri et al., 2021; Dangol et al., 2022). Among the 15 CYPs identified by co-expression analysis, we selected two root-enriched CYP candidates, *CYP1* and *CYP2*, annotated as putative tryptamine 5-hydroxylases (T5Hs), for functional characterization as potential TAM **3** hydroxylases (**Supplemental Table S5**). The two candidates share 79.3% amino acid sequence identity and exhibit 51.48% (*CYP1*) and 53.86% (*CYP2*) identity to the characterized rice T5H (OsCYP71P1, LOC_Os12g16720). Phylogenetic analysis placed both candidates within a clade containing functionally characterized plant T5Hs, further supporting their predicted roles as TAM **3** hydroxylases (**Supplemental Figure S7** and **Supplemental Table S6**). We transiently co-expressed *PhaTDC1* with either *CYP1* or *CYP2* in *N. benthamiana*, and detected the production of 6-HT **6** and 5-HT **9**, respectively (**Figure 2D**). These results establish the distinct regioselectivities of the two CYPs, with CYP1 (CYP71BG47, PhaT6H) and CYP2 (CYP71BG48, PhaT5H) catalyzing TAM **3** hydroxylation at the C6 or C5 positions, respectively (**Figure 2D**). These findings support the observed accumulation of 5-HT **9** in *P. harmala* (**Figure 1A**) and involvement of 6-HT **6** in the early steps of β-carboline biosynthesis. Unexpectedly, co-expression of *PhaTDC1* and *PhaT6H* also resulted in the accumulation of harmalol **7** (**Figure 2E**), suggesting that endogenous host enzymes and/or nonenzymatic reactions contribute to β-carboline scaffold formation from 6-HT **6** in *N. benthamiana* (**Figure 2F**).

### Pyruvate decarboxylation precedes β-carboline scaffold formation

Given the potential role of acetaldehyde **12** in Pictet-Spengler condensation with 6-HT **6** (**Supplemental Figure S1D**), we next investigated whether pyruvate **4** decarboxylation contributes to the accumulation of harmalol **7** and related intermediates. We transiently co-expressed each of the three identified PDC candidates (*PDC1*-*PDC3,* **Supplemental Table S5**) with *PhaTDC1* and *PhaT6H* in *N. benthamiana*. Although none significantly altered harmalol **7** accumulation (**Supplemental Figure S8A**), co-expression of *PDC1* (*PhaPDC1*) resulted in the production of THH-OH **15** (**Supplemental Figure S8B**), a tetrahydro-β-carboline previously detected in *B. caapi* tissues (**Figure 1A** and **Supplemental Figure S4A** and **S8B**), consistent with its generation through a Pictet-Spengler-type reaction between 6-HT **6** and acetaldehyde **12** (**Supplemental Figure S1D**). Neither of the other two PDC candidates supported THH-OH **15** accumulation (**Supplemental Figure S8B**). To directly evaluate the catalytic activity of PhaPDC1, we next transiently expressed *PhaPDC1* or *CmPDC1*, a functionally characterized PDC from melon (*Cucumis melo* L.; MELO3C009145P1, Melonomics) (Wang et al., 2019), in *N. benthamiana* leaves, supplementing the infiltrated tissues with pyruvate **4** as the substrate. Using gas chromatography-mass spectrometry (GC-MS), we detected a significant increase in acetaldehyde **12** in leaves expressing either PDC compared with leaves infiltrated with an empty vector (**Figure 3A**). Transient co-expression of *CmPDC1* with *PhaTDC1* and *PhaT6H* also resulted in the accumulation of THH-OH **15**(**Figure 3B**). Notably, the levels of acetaldehyde **12** were reduced in *N. benthamiana* leaves co-expressing *PhaPDC1*, *PhaTDC1* and *PhaT6H* compared with leaves expressing *PhaPDC1* alone (**Figure 3A**). Together, the ability of PDCs from two different plant species to support THH-OH **15** accumulation, coupled with the depletion of acetaldehyde **12** upon co-expression of *PhaPDC1* with the identified enzymes, supports a role for PDC-derived acetaldehyde **12** as the carbonyl donor in β-carboline scaffold formation (**Figure 3A** and **3B**, and **Supplemental Figure S8B**).

**Figure 3.**
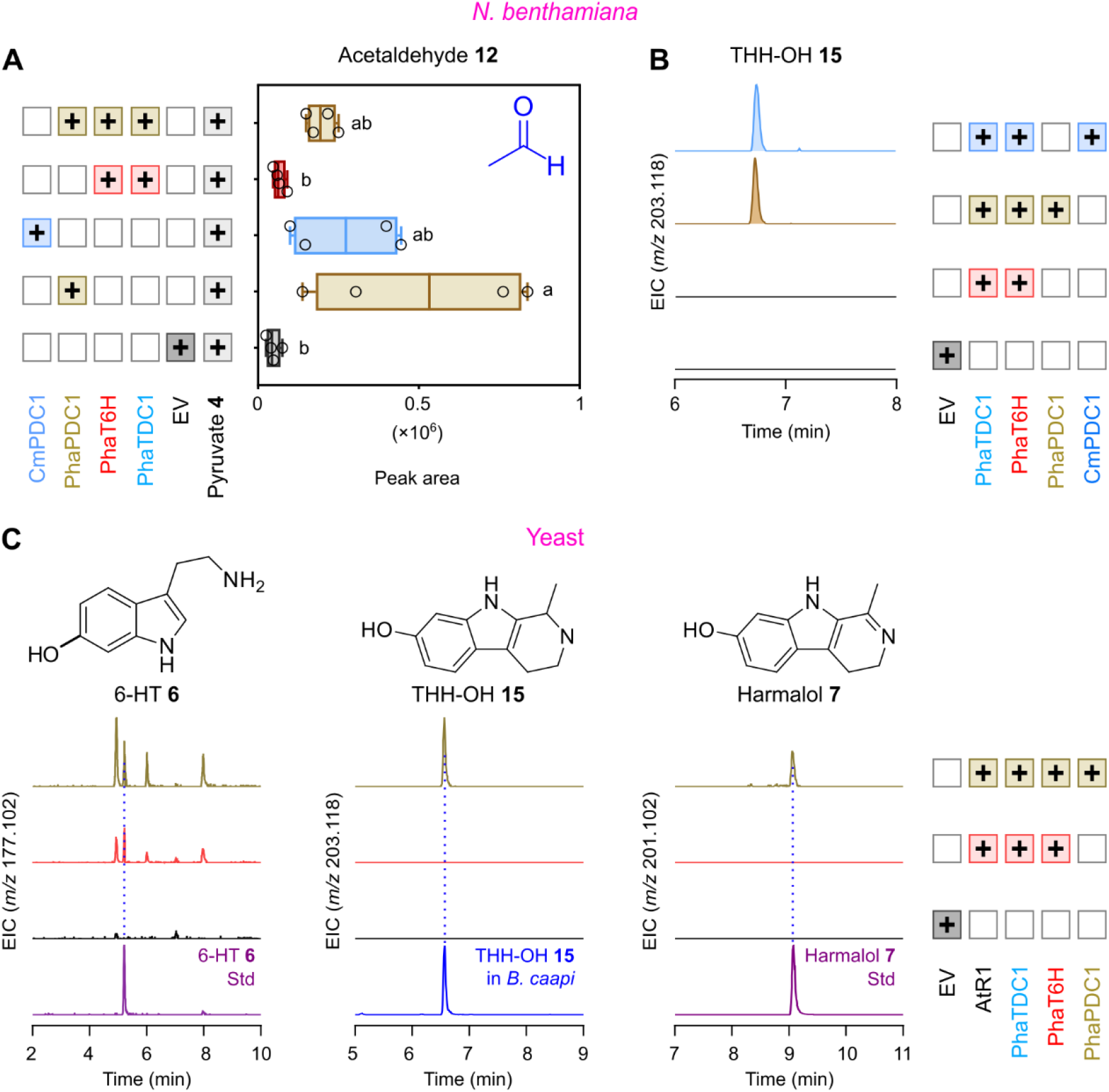
Functional characterization of PhaPDC1 reveals an acetaldehyde-dependent route to β-carboline scaffold formation. **(A)** Ion abundance peak areas (median ± SD; *n* = 4 biological replicates) of acetaldehyde **12** analyzed by gas chromatography-mass spectrometry (GC-MS) following transient co-expression of *PhaTDC1*, *PhaT6H*, and *PhaPDC1* or *CmPDC1* (pyruvate decarboxylase from *Cucumis melo*) in *Nicotiana benthamiana* leaves. Pyruvate **4** was supplied two days after agroinfiltration, and samples were harvested 5 days after infiltration. Statistical significance was assessed by one-way ANOVA followed by Tukey’s HSD test; letters denote statistically significant differences (*p* < 0.05). Acetaldehyde **12** was identified by matching its mass spectral fragmentation pattern to reference spectra in the NIST library. **(B)** Extracted ion chromatograms (EICs) of tetrahydroharmol (THH-OH **15**) following transient expression of *PhaPDC1* or *CmPDC1* together with *PhaTDC1* and *PhaT6H*. The continuation of Figure 3 appears on the next page. **(C)** EICs of 6-hydroxy-tryptamine (6-HT **6**), THH-OH **15**, and harmalol **7** following heterologous co-expression of *PhaTDC1* and *PhaT6H*, with or without *PhaPDC1*, together with *AtR1* in *Saccharomyces cerevisiae* (*n* = 4 biological replicates). Chromatograms were normalized to the highest values. All metabolites were identified using authentic analytical standards, except for THH-OH **15**, which was tentatively identified by comparison of its retention time and MS/MS fragmentation pattern with those of the corresponding metabolite detected in a reference extract of *Banisteriopsis caapi* (**Supplemental Figure S8C**). Plus signs (+) indicate biosynthetic genes expressed in each experiment, with boxes colored according to the last enzyme added in a set of coexpressed genes. Corresponding chemical structures are included for reference. EV, empty vector; Std, standard.

We further co-expressed *PhaTDC1* and *PhaT6H*, with or without *PhaPDC1,* in *Saccharomyces cerevisiae* (*S. cerevisiae*), together with the *Arabidopsis thaliana* NADPH-CYP reductase *AtR1* to support plant CYP activity in yeast (Urban et al., 1997). Expression of *PhaTDC1* and *PhaT6H* resulted in accumulation of 6-HT **6**, confirming TAM 6-hydroxylation activity by *PhaT6H* detected in *N. benthamiana* leaves (**Figure 3C** and **Supplemental Figure S8C**). Notably, THH-OH **15** and harmalol **7** accumulated in yeast only when *PhaPDC1* was co-expressed, consistent with the results obtained in *N. benthamiana* (**Figure 3C** and **Supplemental Figure S8C**). These findings support a role for PDC-derived acetaldehyde **12** in β-carboline scaffold formation.

We next tested several proteins implicated in Pictet-Spengler-type or related condensation reactions (Lichman, 2021) for their potential to catalyze β-carboline scaffold formation. Three candidates were selected based on sequence similarity to members of the Bet v 1-like (*PS1*), strictosidine synthase-like (*PS2*), and major latex protein-like (MLP-like, *PS3*) families (Maresh et al., 2008; Roddan et al., 2020; Singh et al., 2018) (**Supplemental Table S5**). However, transient co-expression of these candidates in *N. benthamiana* leaves together with *PhaTDC1, PhaT6H*, and *PhaPDC1* did not result in increased accumulation of 6-HT **6**, harmalol **7**, or THH-OH **15** (**Supplemental Figure S8D**).

To investigate whether the β-carboline backbone of THH-OH **15** can be generated through nonenzymatic Pictet-Spengler-type reactions, we performed cell-free assays in which pyruvate **4**, acetaldehyde **12**, and 6-HT **6** were incubated individually or in combination at pH 7.4 and 37 °C for 1 h, following the conditions described in (Callaway et al., 1994). We detected THH-OH **15** peaks in reactions containing both acetaldehyde **12** and 6-HT **6** (**Supplemental Figure S9A**), while reactions of pyruvate **4** and 6-HT **6** resulted in harmalol **7** (**Supplemental Figure S9B**). Interestingly, we detected a previously unreported peak with an accurate mass consistent with that of 6-hydroxy-1,2,3,4-tetrahydro-β-carboline-1-carboxylic acid (6-H-THβC-1-COOH **17**) in reactions containing pyruvate **4** and 6-HT **6 (Supplemental Figure S9C** and **S9D**). The detection of this putative intermediate in the cell-free reactions is consistent with a reaction proceeding *via* a carboxylated tetrahydro-β-carboline intermediate followed by its spontaneous oxidative decarboxylation to harmalol **7** (**Supplemental Figure S9E**). Together, our findings provide biochemical evidence that 6-HT **6** can undergo nonenzymatic scaffold-forming reactions with either pyruvate **4** or acetaldehyde **12**, supporting the two proposed routes for β-carboline scaffold formation (**Supplemental Figure S1D**).

### A single OMT catalyzes multiple methylation steps in β-carboline biosynthesis

To identify the active OMTs, we selected four genes from the 15 OMT transcripts identified in the *PhaTDC1*-centered co-expression network based on their expression patterns and sequence similarity to characterized plant OMTs. The four candidates (*OMT1-4*, **Supplemental Table S5**) were recombinantly expressed in *E. coli*, and their activities were evaluated in *in vitro* assays using the corresponding cell lysates, S-Adenosyl-L-methionine (SAM) as the methyl donor, and either harmalol **7** or harmol **8** as substrates. Only OMT2 (PhaOMT2) catalyzed the conversion of harmalol **7** to harmaline **5** and harmol **8** to harmine **1**, whereas the other candidates showed no detectable activity (**Figure 4A**). We next purified the recombinant protein and confirmed its dual-substrate activity (**Supplemental Figure S10A** and **S10B**). Transient expression of *PhaOMT2* in *N. benthamiana* leaves by agro-infiltration, followed two days later by separate supplementation with harmalol **7** or harmol **8**, resulted in accumulation of harmaline **5** and harmine **1**, respectively (**Figure 4B**). Neither product was detected in the corresponding controls, suggesting that *N. benthamiana* lacks endogenous OMT activity capable of catalyzing these reactions. To test whether PhaOMT2 could also *O*-methylate 6-HT **6**, we co-expressed *PhaTDC1*, *PhaT6H*, and *PhaPDC1* with or without *PhaOMT2*. However, 6-MeO-TAM **16** was not detected in any of these combinations (**Supplemental Figure S10C**), consistent with its absence from the metabolite profiles of *P. harmala* (**Figure 1A**). On the other hand, we observed PhaOMT2-mediated *O*-methylation of THH-OH **15** to THH **14**, providing additional support for the tentative structural assignment of THH-OH **15** (**Supplemental Figure S10D**). Interestingly, replacing *PhaPDC1* with *CmPDC1* did not result in detectable THH **14**, although both enzymes supported THH-OH **15** production (**Supplemental Figure S10D**). These results support a role for PhaOMT2 in β-carboline biosynthesis, demonstrating its selectivity for substrates containing the β-carboline scaffold over non-cyclized tryptamines.

**Figure 4.**
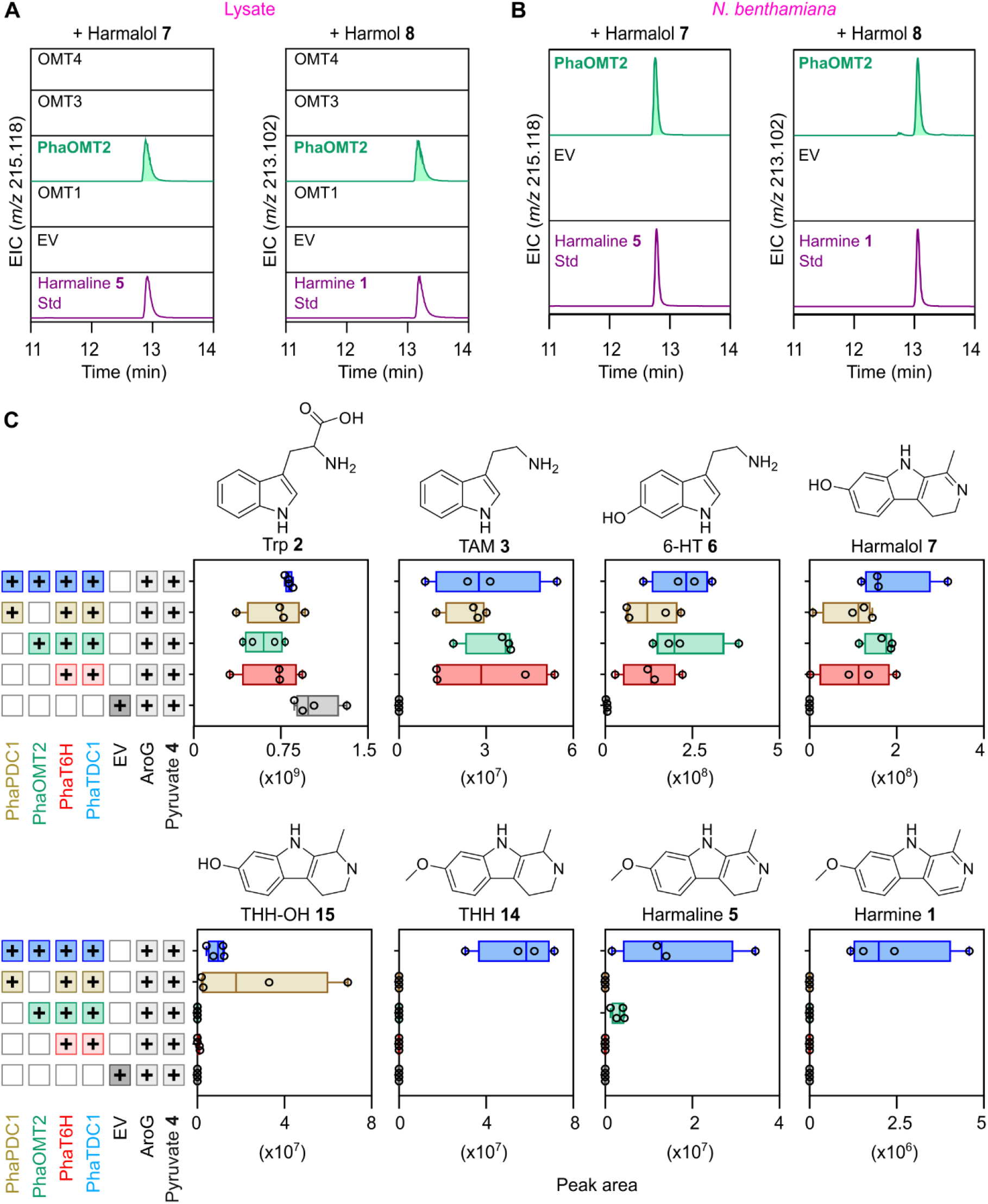
Functional characterization of PhaOMT2 enables heterologous reconstruction of harmine biosynthesis in *Nicotiana benthamiana*. Extracted ion chromatograms of harmaline 5 and harmine 1 following (A) *in vitro* enzyme assays using *Escherichia coli* (*E. coli*) lysates expressing individually OMT1-OMT4 proteins, with S-Adenosyl methionine (SAM) and harmalol 7 or harmol 8 as substrates (*n* = 3 biological replicates), or (B) transient expression of *PhaOMT2* in *Nicotiana benthamiana* (*N. benthamiana*) leaves, (*n* = 4 biological replicates). The continuation of Figure 4 appears on the next page. Leaves were supplemented with harmalol **7** or harmol **8** as substrates 2 days after agroinfiltration and harvested 3 days later. Chromatograms were normalized to the highest values. **(C)** Ion abundance peak areas of β-carbolines and intermediates detected following transient expression of different combinations of *PhaTDC1*, *PhaT6H*, *PhaPDC1*, and *PhaOMT2* in *N. benthamiana* leaves together with the bacterial *AroG* mutant which increases the accumulation of aromatic amino acids in *N. benthamiana* leaves (Berman et al., 2026). Pyruvate **4** was supplied 2 days after agroinfiltration, and samples were harvested 3-5 days later. Data are presented as median ± SD (*n* = 4 biological replicates). Plus signs (+) indicate biosynthetic genes expressed in each experiment, with boxes colored according to the last enzyme added in a set of coexpressed genes. Corresponding chemical structures are included for reference. All metabolites were identified using authentic analytical standards, except for THH-OH **15**, which was tentatively identified by comparison of its retention time and MS/MS fragmentation pattern with those of the corresponding metabolite detected in a reference extract of *Banisteriopsis caapi* (**Supplemental Figure S11**). Trp **2**, tryptophan; TAM **3**, tryptamine; 5-HT **9**, 5-hydroxy-tryptamine; 6-HT **6**, 6-hydroxy-tryptamine; THH **14**, tetrahydroharmine; EV, empty vector; Std, standard.

### Reconstruction of β-carboline biosynthesis in *N. benthamiana*

Sequential introduction of *PhaTDC1*, *PhaT6H*, *PhaPDC1,* and *PhaOMT2* together with a feedback-insensitive mutant of bacterial deoxy-D-arabino-heptulosonate 7-phosphate synthase, to enhance Trp **2** levels (Tzin et al., 2013; Berman et al., 2026) resulted in accumulation of TAM **3**, 6-HT **6**, harmalol **7**, THH-OH **15**, THH **14**, and unexpectedly, also harmaline **5**, and harmine **1** (**Figure 4C** and **Supplemental Figure S11**). Notably, THH **14** was only observed upon co-expression of *PhaPDC1* and *PhaOMT2*, which also increased the accumulation of harmaline **5** and harmine **1** (**Figure 4C** and **Supplemental Figure S11**). These findings demonstrate that β-carboline biosynthesis can be reconstructed in *N. benthamiana* without introducing a dedicated oxidase, suggesting that spontaneous oxidation or endogenous host enzymes can support the remaining oxidation steps.

To investigate whether *P. harmala* encodes oxidases catalyzing the remaining steps of β-carboline biosynthesis, we focused on FAD-dependent oxidases of the berberine bridge enzyme-like (BBE-like) family, which mediate diverse oxidative reactions in plant specialized metabolism, including the oxidative dehydrogenation of reduced alkaloids in *Papaver somniferum* (Hagel et al., 2012; Bakam et al., 2026). Six BBE-like candidates were selected based on their high expression in roots and co-expression with *PhaTDC1* (**Supplemental Table S5**). In addition, one YUCCA (YUC)-like flavin monooxygenase was also selected for functional testing based on its potential role in oxidative reactions (Zhao et al., 2001). The BBE-like candidates were tested with and without their predicted N-terminal signal peptides to assess enzymatic activity independently of subcellular targeting. Oxidase candidates were transiently co-expressed with *AroG, PhaTDC1*, *PhaT6H*, *PhaPDC1*, and *PhaOMT2* in *N. benthamiana* leaves. However, no increased accumulation of harmalol **7**, THH-OH **15**, THH **14**, harmaline **5,** or harmine **1** was observed under the conditions examined (**Supplemental Figure S12A** and **S12B**).

### Two active-site residues determine CYP regioselectivity for 5-HT and 6-HT production

To understand the molecular basis of the distinct TAM **3** hydroxylation regioselectivities exhibited by the closely related CYPs PhaT6H and PhaT5H, we used AlphaFold3 (Abramson et al., 2024) to model each CYP protein in complex with its respective product, 6-HT **6** or 5-HT **9**, and superimposed the predicted structures (**Figure 5A**). The predicted active sites of the two CYPs were nearly identical, differing at only two positions: Leu373 and Thr490 in PhaT6H, corresponding to Val371 and Ala488 in PhaT5H (**Figure 5A** and **Supplemental Figure S13A**). We hypothesized that the bulkier Leu373 residue in PhaT6H creates steric constraints that disfavor hydroxylation at the C5 position of TAM **3**, thereby favoring C6 hydroxylation. To determine whether these residues govern hydroxylation regioselectivity, we generated single and double mutants of each enzyme (**Figure 5B**). The single mutants PhaT6H(L373V) and PhaT5H(V371L) produced both 6-HT **6** and 5-HT **9**, whereas PhaT6H(T490A) and PhaT5H(A488T) largely retained the products of their respective wild-type enzymes (**Figure 5B**). Remarkably, the double mutants PhaT6H(L373V/T490A) and PhaT5H(V371L/A488T) completely reversed their hydroxylation regioselectivity (**Figure 5B**). AlphaFold3 modeling further revealed that the active site of the rice T5H (OsCYP71P1) closely resembles that of PhaT5H (**Supplemental Figure S13B**). Together, these findings identify two key residues governing the regioselectivity of TAM **3** hydroxylation and suggest that changes in substrate binding orientation within the CYP active site may have facilitated the emergence of 6-HT **6** biosynthesis in *P. harmala* and its recruitment into the β-carboline pathway.

**Figure 5.**
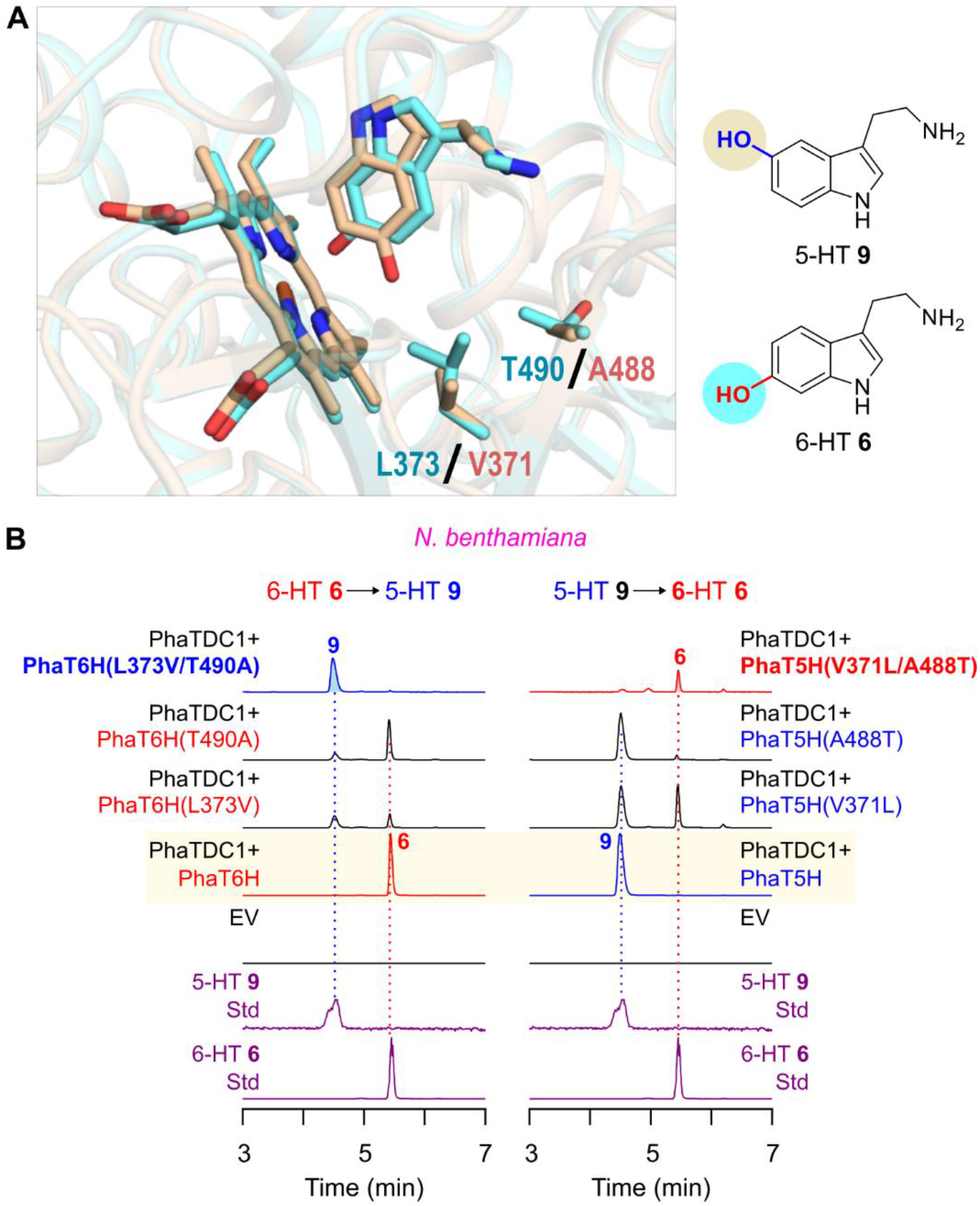
Structural and functional characterization of hydroxylation regioselectivity in *Peganum harmala* cytochrome P450 (CYP) enzymes. (A) Overlay of AlphaFold3-predicted structures of PhaT6H in complex with 6-hydroxy-tryptamine (6-HT 6, cyan) and PhaT5H in complex with 5-hydroxy-tryptamine (5-HT 9, wheat). The heme groups, bound ligands, and two differing active-site residues are shown as sticks. Leu373 and Thr490 in PhaT6H, corresponding to Val371 and Ala488 in PhaT5H, are proposed to influence substrate positioning and hydroxylation regioselectivity. (B) Extracted ion chromatograms of 5-HT 9 and 6-HT 6 following transient co-expression of *PhaTDC1* with wild-type *PhaT6H* or *PhaT5H* (highlighted in yellow), or their respective mutants in *Nicotiana benthamiana* leaves (*n* = 4 biological replicates). Amino acid substitutions are indicated for each mutant. The continuation of Figure 5 appears on the next page. Samples were collected four days after infiltration. 6-HT **6** and 5-HT **9** were identified using analytical standards. Chromatograms were normalized to the highest values. EV, empty vector; Std, standard.

### Engineering ayahuasca-associated alkaloid biosynthesis in a single plant system

We next explored whether the biosynthetic pathways of DMT **13** and harmine **1**, which share TAM **3** as a common precursor but occur naturally in different plant species **Supplemental Figure S14A**), could be functionally integrated within a single plant system. We employed two complementary approaches: transient co-expression of both pathways in *N. benthamiana,* and engineering of the DMT **13** pathway into β-carboline-producing *P. harmala* hairy roots. In the first approach, we transiently co-expressed *PhaTDC1*, *PhaT6H*, *PhaPDC1*, and *PhaOMT2* for β-carboline biosynthesis, together with the previously characterized DMT **13** biosynthetic genes from *P. viridis*, *PvTDC2* and *PvNMT1* (Berman et al., 2026), using *AroG* to enhance precursor availability, in *N. benthamiana* leaves. This resulted in simultaneous accumulation of harmine **1**, DMT **13**, and their biosynthetic intermediates in the same tissue (**Figure 6A** and **Supplemental Figure S14B**).

**Figure 6.**
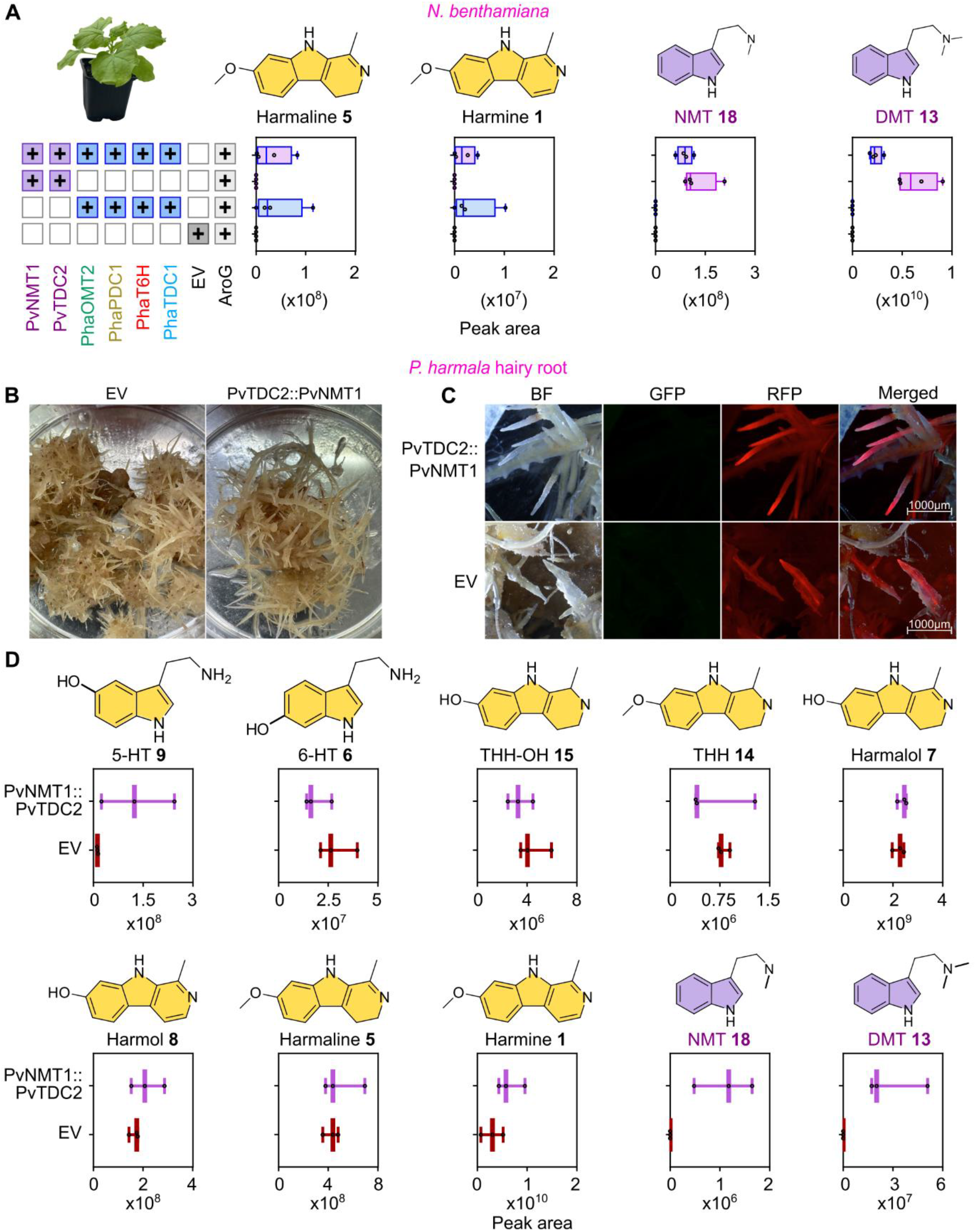
Combined production of β-carbolines and *N*,*N*-dimethyltryptamine. (**DMT) in a single plant. (A)** Representative image of *Nicotiana benthamiana* (*N. benthamiana*), ion abundance peak areas of harmine **1**, DMT **13**, and their biosynthetic intermediates following transient expression of harmine **1** biosynthetic genes (*PhaTDC1*, *PhaT6H*, *PhaPDC1*, and *PhaOMT2*), DMT **13** biosynthetic genes (*PvTDC2* and *PvNMT1*), or their combination. The continuation of Figure 6 appears on the next page. The bacterial *AroG* mutant was co-expressed in all combinations to enhance the accumulation of aromatic amino acids in *N. benthamiana* leaves. Data are presented as median ± SD (*n* = 4 biological replicates). Samples were harvested seven days after infiltration. Plus signs (+) indicate biosynthetic genes expressed in each experiment, with blue, purple, and combined blue-purple boxes representing β-carboline, DMT **13**, and combined biosynthetic genes, respectively. **(B-C)** Representative bright-field (BF), green fluorescence protein (GFP) or red fluorescence protein (RFP) reporters, and merged images of transformed *Peganum harmala* (*P. harmala*) hairy roots generated from *P. harmala* cotyledons following *Rhizobium rhizogenes* (MSU440)-mediated transformation with a construct expressing *PvTDC2* and *PvNMT1* from *Psychotria viridis* (Berman et al., 2026), and RFP as a reporter. **(D)** Ion abundance peak areas (median ± SD; *n* = 3 biological replicates) of harmine **1**, DMT **13**, and their intermediates detected in transgenic hairy roots. Hairy roots transformed with an RFP-expressing empty vector (EV) served as the control. All metabolites were identified using authentic analytical standards, except for THH-OH **15**, which was tentatively identified by comparison of its retention time and MS/MS fragmentation pattern with those of the corresponding metabolite detected in a reference extract of *Banisteriopsis caapi*. Corresponding chemical structures are included for reference. 5-HT **9**, 5-hydroxy-tryptamine; 6-HT **6**, 6-hydroxy-tryptamine; THH **14**, tetrahydroharmine; NMT **18**, *N*-methyltryptamine.

Next, we generated transgenic *P. harmala* hairy roots through *Agrobacterium rhizogenes*-mediated transformation of cotyledons with constructs expressing *PvTDC2* and *PvNMT1*, followed by selection of transformed tissues based on RFP fluorescence. RFP-positive calli developed and subsequently formed transgenic hairy roots within approximately two months (**Figure 6B, 6C**, and **Supplemental Figure S15A**). Similar to the reconstruction in *N. benthamiana*, transformed *P. harmala* tissues simultaneously accumulated harmine **1** and DMT **13**, together with their respective biosynthetic intermediates, whereas control tissues expressing RFP alone exhibited comparable β-carboline profiles but no detectable NMT **18** or DMT **13** (**Figure 6D** and **Supplemental Figure S15B** and **S15C**). Notably, THH-OH **15** and THH **14**, which were not detected in native *P. harmala* roots (**Figure 1A**), accumulated in transformed calli and hairy roots (**Figure 6D** and **Supplemental Figure S15B**), potentially reflecting altered metabolic flux or reduced turnover of these intermediates in transformed tissues. Their accumulation further supports their proposed participation in the biosynthetic pathway, consistent with our functional assays. Together, the simultaneous production of β-carbolines and DMT **13** reconstitutes the characteristic combination of β-carbolines and psychedelic tryptamines associated with ayahuasca brew within a single plant system and demonstrates the feasibility of integrating distinct alkaloid biosynthetic pathways through metabolic engineering.

## Discussion

The reported antimicrobial, antifungal, and anti-herbivore activities of β-carboline alkaloids suggest that these metabolites may serve diverse ecological functions. The spatial distribution of these metabolites may provide further insights into these functions, as their accumulation in tissues exposed to the surrounding environment could facilitate interactions with soil- and surface-associated organisms. In this context, the localization of harmine **1** in the epidermal regions of *P. harmala* roots and outer stem regions of *B. caapi* may support a role in chemical defense. The preferential accumulation of β-carbolines in the roots of *P. harmala*, compared with their broader distribution across tissues in *B. caapi*, may reflect differences in ecological adaptation between these species, as *P. harmala* inhabits arid and semi-arid environments, whereas *B. caapi* is a tropical Amazonian liana.

Based on our findings, we propose a biosynthetic pathway for β-carbolines in *P. harmala*, beginning with PhaTDC1-mediated decarboxylation of Trp **2** to TAM **3**, followed by PhaT6H-catalyzed hydroxylation to 6-HT **6** (**Figure 7**). The identification of PhaT6H establishes a previously unresolved enzymatic link in β-carboline biosynthesis. The distinct regioselectivities of PhaT6H and its closely related homolog PhaT5H illustrate how limited active-site divergence may enable the emergence of alternative metabolic pathways by directing shared precursors toward chemically distinct products. The competing 5-hydroxylation branch may also influence precursor availability for β-carboline biosynthesis, consistent with previous feeding experiments in *P. harmala* hairy roots, in which exogenous TAM **3** preferentially increased 5-HT **9** rather than β-carboline production (Berlin et al., 1993).

**Figure 7.**
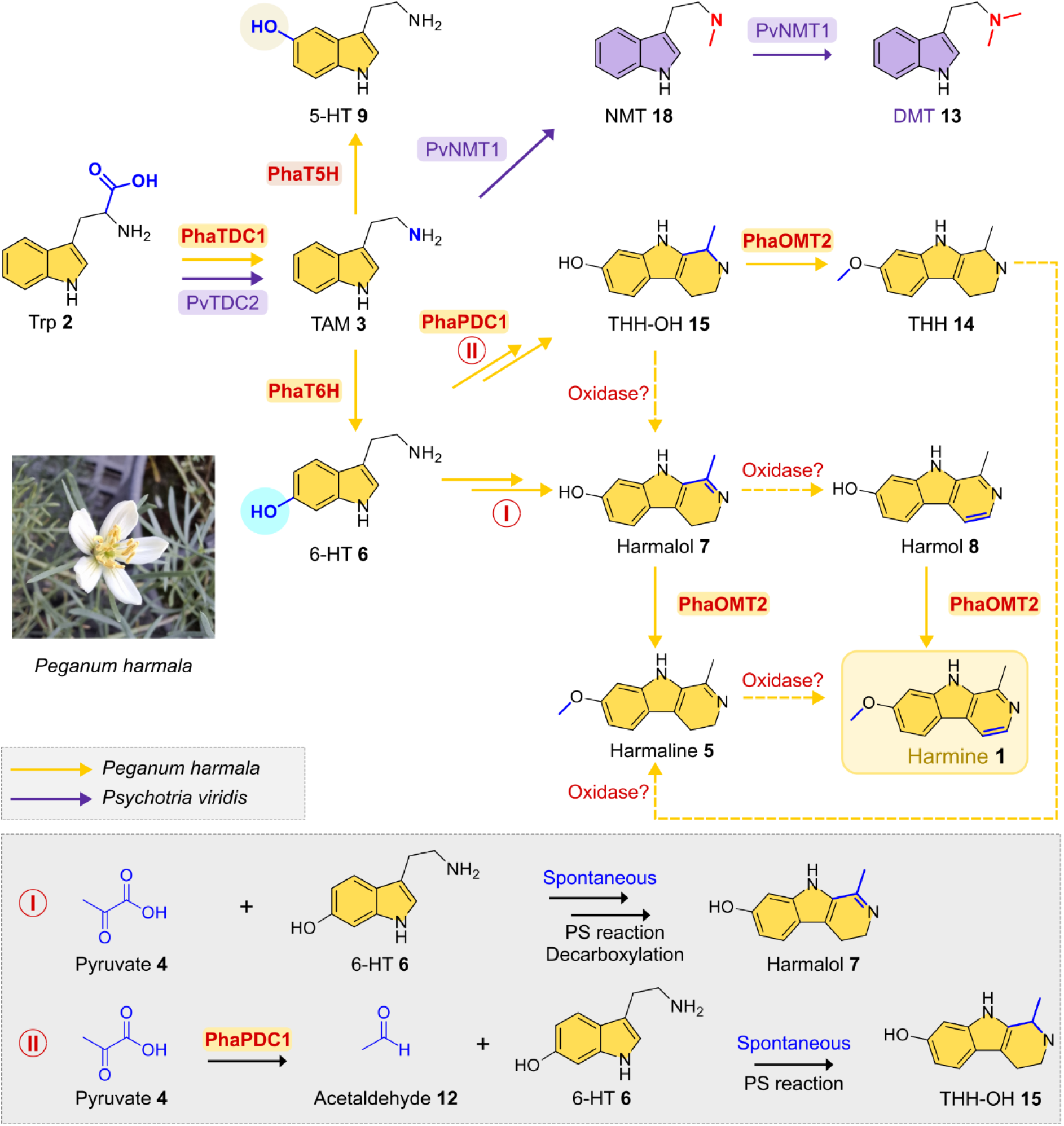
Proposed β-carboline biosynthetic pathway in *Peganum harmala*. Modules I and II illustrate two proposed routes for β-carboline scaffold formation through Pictet-Spengler (PS) reactions. Yellow arrows indicate the proposed biosynthetic pathway in *Peganum harmala* (*P. harmala*), and dashed arrows experimentally unconfirmed steps. Enzymes functionally characterized from *P. harmala* are highlighted in yellow or red. Enzyme classes marked with question marks indicate reactions for which specific enzymes remain unidentified. The hydroxyl groups of 6-hydroxytryptamine (6-HT **6**) and 5-hydroxytryptamine (5-HT **9**) are highlighted in cyan and wheat, respectively, to indicate their distinct hydroxylation positions. The continuation of Figure 7 appears on the next page. The *N*,*N*-dimethyltryptamine (DMT **13**) biosynthetic pathway discovered previously in Psychotria viridis (Berman et al., 2026), including its enzymes, metabolites, and reaction arrows, is shown in purple to illustrate its biosynthetic relationship with harmine **1**. Trp **2**, tryptophan; TAM **3**, tryptamine; THH-OH **15**, tetrahydroharmol; THH **14**, tetrahydroharmine; NMT **18**, *N*-methyltryptamine.

Accordingly, suppressing PhaT5H activity in this system may provide a strategy to redirect metabolic flux toward increased β-carboline production. More broadly, our findings highlight the potential of AlphaFold 3-guided structural modeling and protein design to elucidate how specific active-site residues influence substrate binding and positioning, thereby determining enzyme specificity and regioselectivity, and guide protein engineering to tailor enzyme activity. Several routes have been proposed for the biosynthetic assembly of the β-carboline scaffold, but their biochemical basis has remained unresolved (**Supplemental Figure S1**). Our findings support a pathway in which 6-HT **6**, rather than TAM **3**, undergoes Pictet-Spengler-type cyclization through two alternative routes: a putative nonenzymatic Pictet-Spengler reaction between 6-HT **6** and pyruvate **4**, followed by oxidative decarboxylation to harmalol **7** (Module I in **Figure 7**), or a nonenzymatic Pictet-Spengler condensation of 6-HT **6** with acetaldehyde **12**, generated through PhaPDC1-mediated decarboxylation of pyruvate **4**, to yield THH-OH **15** (Module II in **Figure 7**). Subsequent oxidation steps may occur spontaneously or involve endogenous enzymes that remain to be characterized. The spontaneous formation of β-carboline intermediates demonstrates the chemical feasibility of scaffold assembly without a dedicated PS enzyme. However, spontaneous and enzyme-catalyzed reactions may occur in parallel, and the contribution of endogenous enzymes, including potentially promiscuous PS enzymes, to these reactions cannot be excluded. This raises the possibility that the emergence of β-carboline biosynthetic pathways may depend on the evolution of enzymes that generate suitable reactive intermediates, without requiring dedicated catalysts for every subsequent transformation. The activity of PhaOMT2 toward multiple β-carboline intermediates, including harmalol **7**, harmol **8**, and THH-OH **15**, provides another example of this biosynthetic flexibility (**Figure 7**).

The involvement of PhaPDC1 in this pathway reveals an unexpected role for an enzyme primarily associated with central carbon metabolism in specialized alkaloid biosynthesis. In addition to the established functions of PDCs in fermentation, the pyruvate dehydrogenase bypass, and volatile aroma biosynthesis (Wang et al., 2019), our findings demonstrate that PDC-derived acetaldehyde **12** can serve as a reactive precursor for spontaneous PS-type condensation with 6-HT **6**. To our knowledge, this is the first example of PDC-derived acetaldehyde 12 serving as a reactive precursor for alkaloid biosynthesis in plants. Notably, PhaPDC1 was predicted to localize to plastids based on the presence of a chloroplast transit peptide (localization probability of 0.97; **Supplemental Table S5**), whereas CmPDC1 was previously reported to localize to the cytosol (Wang et al., 2019). Despite their predicted differences in subcellular localization, both enzymes supported THH-OH **15** accumulation in *N. benthamiana*, whereas only PhaPDC1 showed detectable THH **14** accumulation. These differences suggest that precursor availability, subcellular compartmentalization, and downstream metabolic activities may influence the relative contributions of alternative biosynthetic routes and the resulting metabolic profiles.

The complementary engineering of *N. benthamiana* and *P. harmala* hairy roots demonstrates two strategies for the production of high-value TAM **3**-derived alkaloids. Transient expression in *N. benthamiana* provides a flexible platform for reconstructing and combining heterologous pathways, paving the way for their stable introduction into engineered plants, either individually or in combination. In contrast, *P. harmala* hairy roots enable the introduction of new biosynthetic capabilities into a native β-carboline-producing system, where targeted silencing of competing metabolic branches may enhance product accumulation and facilitate the discovery of additional biosynthetic enzymes, transporters, and other pathway-associated factors. This platform could be further expanded by incorporating additional TAM **3**-derived pathways, building on our previous reconstruction of psychedelic tryptamines and their halogenated derivatives in plants (Berman et al., 2026). Such modular engineering strategies offer opportunities to produce diverse natural and new-to-nature alkaloids while investigating precursor competition, pathway compatibility, and the biochemical factors governing their production.

The growing pharmacological and therapeutic interest in β-carbolines and DMT **13**, together with their reliance on plant-derived sources, highlights the need for sustainable production strategies. Given the importance of β-carbolines and DMT **13** in ayahuasca preparations, stable engineered plants could provide renewable sources of these compounds and their derivatives, supporting further pharmaceutical and biotechnological research. Such approaches could reduce reliance on wild harvesting, particularly of slowly regenerating plants, alleviate pressure on natural populations, and contribute to the conservation of plant genetic diversity and associated ecosystems.

## Methods

### Chemicals and plant material

All analytical standards used in the study were of ≥ 95% purity. Trp **2** and TAM **3** were obtained from Sigma-Aldrich (Rehovot, Israel). 5-MeO-TAM, THβC-1-COOH **10**, 5-HT **9**, 6-HT **6**, THH **14**, harmalol **7**, harmaline **5**, harmol **8**, and harmine **1** were purchased from Cayman Chemical (Ann Arbor, MI, USA). *Peganum harmala* seeds were obtained from the Israel Gene Bank, and *B. caapi* young plants were sourced from a local nursery. Plants were grown in a greenhouse under long-day conditions, with day/night temperatures maintained at 24/18 °C, respectively. Tissues from *P. harmala* and *B. caapi* plants were collected at approximately one year of age for chemical profiling and RNA extraction.

### High resolution liquid chromatography-mass spectrometry (LC-MS) chemical analysis

Unless otherwise stated, 100 mg of frozen powdered plant tissue was extracted with 300 µL 80% methanol containing 0.1% formic acid. Samples were sonicated for 30 min, agitated for an additional 30 min, and centrifuged at 14,000 g for 10 min. The resulting supernatants were filtered through 0.22 µm syringe filters prior to LC-MS/MS analysis. Metabolite profiling was performed using an ultra-performance liquid chromatography system (UPLC; Waters Acquity) equipped with a diode array detector coupled to either an Orbitrap Exploris 240 MS (Thermo Scientific, Bremen, Germany) or an Orbitrap IQ-X Tribrid mass spectrometer. Chromatographic separation was achieved using a Waters Acquity UPLC HSS-T3 C18 column (100 mm × 2.1 mm i.d., 1.7 µm particle size). The mobile phases consisted of (A) 0.1% formic acid in acetonitrile: water (1:99, v/v) and (B) 0.1% formic acid in acetonitrile. The flow rate was maintained at 0.3 mL min^−1^, and the column temperature was set to 35 °C. The gradient program was as follows: 0% B for 2 min, increased to 75% B over 14 min, raised to 100% B within 0.2 min, maintained at 100% B until 19 min, returned to 0% B within 0.2 min, and equilibrated the system at 0% B until 22 min. Mass spectrometry was performed using electrospray ionization (ESI) in positive ionization mode over an *m/z* range of 70-1,000. Source parameters were set as follows: sheath gas, auxiliary gas, and sweep gas flow rates of 45, 10, and 1 arbitrary units, respectively; vaporizer and ion transfer tube temperatures of 300 °C; and spray voltage of 3.5 kV. Data were acquired in full MS^1^ mode at 60,000 resolution, followed by data-dependent MS/MS (dd-MS^2^) acquisition at 15,000 resolution. Both acquisition modes used standard normalized automatic gain control (AGC) targets and automatic maximum injection times. The dd-MS^2^ settings included a 1.5 m/z isolation window and stepped normalized collision energies of 30, 45, and 60. Compounds with analytical standards were identified by comparison of their retention times and MS/MS spectra with those of the corresponding standards, whereas compounds without available analytical standards were tentatively annotated based on accurate mass, relative retention time, and MS/MS fragmentation patterns (**Supplemental Figure S2** and **S10**). Since minor differences in fragmentation spectra, retention times, and ion abundance were observed between instruments, metabolites were confirmed by comparison with analytical standards and/or extracts analyzed under matched experimental conditions.

### Absolute quantification of β-carbolines

Fresh bark, pith, sap, stem, leaf, and root tissues were collected from *B. caapi*, and young and mature roots, stems, and leaves were collected from *P. harmala*. For metabolite extraction, 100 mg of frozen, powdered tissue was weighed in triplicate and extracted with 1 ml of 80% methanol containing 0.1% formic acid, following the procedure described above. Absolute metabolite quantification was performed using external calibration curves. To ensure that analyte concentrations fell within the linear range of the calibration curves, samples were analyzed at multiple dilutions. LC-MS analyses were conducted on an IQ-X Tribrid mass spectrometer operating in full MS^1^ acquisition mode. In addition to absolute quantification, samples were subjected to comparative metabolite profiling, with peak areas used for relative abundance analyses.

### MALDI-MSI of *B. caapi* stems and *P. harmala* roots

For metabolite localization in *B. caapi* and *P. harmala* tissues, samples were cryo-sectioned (*n*=*2* biological replicates) and coated with matrix using a TM Sprayer (HTX Technologies, USA). The matrix solution consisted of 2,5-dihydroxybenzoic acid (DHB; 40 mg mL^-1^ in 70% containing 0.2% trifluoroacetic acid). Matrix application was performed with the nozzle maintained at 70 °C, using 16 spray passes at a linear velocity of 120 cm min^-1^ and a flow rate of 50 μL min^-1^. Tissue sections were imaged using a Nikon DS-Ri2 microscope. MALDI-MSI was conducted on a 7 T Solarix FT-ICR (Fourier Transform Ion Cyclotron Resonance) mass spectrometer (Bruker Daltonics). Data acquisition and processing were performed using flexImaging v4.1 (Bruker). Spectra were acquired in positive ion mode with lock-mass calibration using the DHB matrix ion ([3DHB+H-3H_2_O]^+^, *m/z* 409.055408). Data were collected at a laser frequency of 1 kHz and 40% laser power, with 200 laser shots per pixel and a spatial resolution of 50 µm. Mass spectra were acquired over an m/z range of 120-2000 in broadband mode with an acquisition time domain of 1M, yielding an estimated resolving power of approximately 115,000 at *m/z* 400. MS/MS spectra were normalized to root-mean-square intensity, and ion images were generated using a mass tolerance of ± 0.005% of the theoretical *m/z* value with pixel interpolation enabled.

### RNA sequencing and annotation of plant transcriptomes

Total RNA was extracted from the same tissue types of *P. harmala* and *B. caapi* used for metabolite profiling, using either the Spectrum Plant total RNA kit (Sigma-Aldrich) or the RNeasy plant mini kit (Qiagen). RNA integrity was assessed using a TapeStation instrument. Extracted RNA from pooled tissues was used for both short-read and long-read transcriptome sequencing. Iso-Seq sequencing libraries for *B. caapi* and *P. harmala* were generated from pooled RNA samples derived from six and four tissues, respectively. Paired-end Illumina libraries were prepared using the TruSeq RNA Library Preparation Kit (Illumina) following the manufacturer’s instructions and sequenced on an Illumina NovaSeq X Plus platform (PE150), generating approximately 40 million reads per sample. Full-length transcript sequencing was performed using the PacBio Iso-Seq workflow. Libraries were prepared using the SMRTbell Prep Kit 3.0 (Pacific Biosciences) from pooled tissue RNA samples of *B. caapi* and *P. harmala* and sequenced on the PacBio Sequel II platform using SMRT Cell 8M and the Sequel II Sequencing Kit, according to the manufacturer’s instructions. Iso-Seq data were processed using the SMRT Link command-line tools. Primer sequences were removed and reads were demultiplexed with lima v2.9.0, using the isoseq and peek-guess switches, and clipped sequences were saved with the dump-clips switch. The isoseq refine v4.0.0 was then used with the require-polya switch to obtain full-length, nonconcatemer (FLNC) reads. FLNC reads were converted to FASTQ with BamTools v2.5.2 and clustered using isONclust v0.0.6.1 (Sahlin and Medvedev, 2020) with the isoseq switch and default parameters (k = 15, w = 50, minimum shared minimizers = 5, minimum average read quality = 7). The representative read of each cluster was extracted into a FASTA file per species, which was used for downstream annotation. Raw Illumina reads were quality-filtered and trimmed using Trimmomatic (Bolger et al., 2014), as implemented in Trinity (Haas et al., 2013) with default parameters. The processed reads were subsequently used for *de novo* transcriptome assembly using Trinity (Haas et al., 2013). Transcript abundances were estimated using RSEM (Li and Dewey, 2011) and normalized using DESeq2 (Love et al., 2014). Functional annotation of predicted transcripts from the Trinity assemblies and the Iso-Seq cluster sequences of both species was performed as described previously (Berman et al., 2024), with additional analyses. Subcellular localization was predicted using TargetP 2.0 (Armenteros et al., 2019) and DeepLoc 2.0 (Thumuluri et al., 2022). Enzymatic functions were predicted with ECPred (Dalkiran et al., 2018) for the Illumina-based transcriptomes and with DeepEC (Ryu et al., 2019) for the Iso-Seq sequences. Completeness of the Iso-Seq transcript sets was assessed with BUSCO v5.7.1 (Manni et al., 2021) in transcriptome mode using the eudicots_odb10 dataset (n = 2,326). Hierarchical clustering was performed using Euclidean distance and complete linkage on trimmed mean of M-values (TMM)-normalized TruSeq transcript expression values converted to counts per million (CPM). Transcripts with CPM > 25 in at least one of 12 sample tissues were retained, resulting 10,905 of 307,954 transcripts (3.54%). Expression values were averaged across three biological replicates for each of four tissues and transformed as log_2_ expression relative to the median expression across tissues. For co-expression analysis, Pearson correlation coefficients were calculated between the bait transcript (*PhaTDC1*) and each isoseq transcript across the analyzed samples. Transcripts with *r* > 0.8 and FDR-adjusted *p* < 0.05 were considered positively co-expressed with *PhaTDC1*. The resulting co-expression network was visualized using Cytoscape (Shannon et al., 2003).

### Sequence alignments and phylogenetic analysis

Protein sequence comparisons were performed using Clustal Omega, and sequence identity was quantified as the proportion of identical residues within the aligned regions. For the phylogenetic analysis of β-carboline-producing plants, amino sequences of maturase K (MatK), encoded by the chloroplast *matK* gene, were retrieved from GenBank. The corresponding protein accession numbers are listed in **Supplemental Table S3**. For the CYP phylogenetic analysis, amino acid sequences of putative *P. harmala* CYPs and functionally characterized plant CYPs were selected from previously published studies of TAM hydroxylases and related enzymes (Fujiwara et al., 2010; Negri et al., 2021; Dangol et al., 2022). Sequences were retrieved from the plant CYP database, and the complete list of protein IDs is provided in **Supplemental Table S5**. A multiple-sequence alignment was generated using MUSCLE, and a maximum-likelihood phylogenetic tree was constructed in MEGA 12 with 1,000 bootstrap replicates. The species-level tree was rooted.

### Recombinant protein expression in *E. coli*

The coding sequence of *PhaTDC1* was amplified by PCR from cDNA of *P. harmala*, and *PhaOMT2* was synthesized by Twist Biosciences. Each gene was individually cloned into the EcoRI-digested pET28a vector using Gibson Assembly (Gibson et al., 2009). All constructs were sequenced-verified and transformed into *E. coli* BL21(DE3) pLysS competent cells. Starter cultures were grown overnight in LB medium supplemented with kanamycin (50 µg mL^-1^) at 37 °C, then diluted 1:100 into fresh LB containing kanamycin and incubated at 37 °C until reaching an optical density at 600 nm (A600) of 0.6. Protein expression was induced with 500 μM isopropyl-1-thio-β-d-galactopyranoside (IPTG), and cultures were further incubated overnight at 16 °C. Cells were harvested by centrifugation at 4,300 rpm for 10 min, and pellets were lysed by sonication in lysis buffer consisting of 50 mM Tris-HCl (pH 8.0), 10% glycerol, 0.1 mg mL^-1^ lysozyme (Sigma-Aldrich), 0.01% benzonase nuclease (Sigma-Aldrich), protease inhibitor cocktail (Sigma-Aldrich), and 0.5 mM phenylmethylsulfonyl fluoride (PMSF, Sigma-Aldrich). The clarified lysate was either used directly for activity assays or subjected to protein purification. Hexahistidine-tagged proteins were purified using a Capturem His-tag purification 24-well plate (Takara Bio, USA) following the manufacturer’s instructions. Elution was performed with 500 mM imidazole in buffer containing 20 mM NaH_2_PO_4_ (pH 7.6) and 500 mM NaCl. Desalting was carried out by repeated centrifugation at 4 °C using Amicon Ultra 10 kDa cutoff filters (Merck Millipore) in Tris-HCl (pH 8.0). Protein purity was evaluated by SDS-PAGE on 10% polyacrylamide gels. Protein purification and enzymatic assays were performed sequentially on the same day.

### TDC and OMT enzyme assays

TDC activity assays were carried out in a total volume of 50 μL containing 100 mM Tris-HCl buffer (pH 8.0), 1 mM pyridoxal-5′-phosphate (PLP), 2 mM Trp **2,** and either 25 μL of crude lysate or 5 μL of purified enzymes. OMT enzyme assays were similarly performed in 50 μL reactions composed of 100 mM Tris-HCl buffer (pH 8.0), 1 mM S-adenosyl-L-methionine (SAM), and 25 μL of crude enzyme lysate. Reactions were incubated at 30 °C for 2 h and terminated by the addition of 70 μL ice-cold methanol with 0.1% formic acid. Samples were vortexed and centrifuged at 13,000 g for 10 min. The supernatants were filtered and analyzed by LC-MS.

### Gas chromatography-mass spectrometry (GC-MS) analysis of acetaldehyde

Leaves of *N. benthamiana* infiltrated with *PhaPDC1* or *CmPDC1*, or with combinations of *PhaTDC1* and *PhaT6H* with or without *PhaPDC1*, were ground into a fine powder, and 1 g of tissue was transferred into GC-MS glass vials containing 1 g of NaCl and 2 mL of 20% NaCl solution (w v^-1^), followed by addition of 20 µL of 2-octanol (5 mg L^-1^) as an internal standard. The vials were sealed with magnetic screw caps filled with silicone/PTFE septa (Thermo Fisher Scientific, Waltham, MA, USA) and shaken for 30 min at 37 °C and 125 rpm. After shaking, acetaldehyde **12** was extracted from the leaf tissue using solid-phase microextraction (SPME) using a 50/30-µm DVB/CAR/PDMS StableFlex fiber (2 cm; Supelco, Bellefonte, PA, USA). GC-MS analysis was performed on an Agilent 7890A gas chromatograph coupled to an Agilent 5977B mass spectrometer using an HP-5MS column (5% phenyl methyl polysiloxane; 30 m x 0.25 mm x 0.25 µm film thickness; Agilent Technologies, Santa Clara, CA, USA). The SPME fiber was desorbed at 250 °C for 5 min in splitless mode, with helium used as the carrier gas. The temperature program was as follows: initial temperature at 40 °C for 1 min; ramp to 90 °C at 5 °C/min; and held at 90 °C for 2 min. The mass spectrometer was operated in electron ionization mode at 70 eV, with an acquisition range of *m/z* 40-280. Acetaldehyde **12** was identified by matching its mass spectral fragmentation pattern to reference spectra in the NIST library, version 14. Four biological replicates from each treatment were analyzed.

### Cell free assays

Cell-free reactions were performed in a total volume of 50 μL containing 100 mM Tris-HCl buffer (pH 7.4) and 1 mM each of pyruvate **4**, acetaldehyde **12**, 6-HT **6**, added individually or in the indicated combinations. Reactions were incubated at 37 °C for 1 h and quenched with 70 μL ice-cold methanol with 0.1% formic acid. Samples were vortexed and centrifuged at 13,000 g for 10 min. The supernatants were filtered and analyzed by LC-MS.

### *Agrobacterium*-mediated transient expression in *N. benthamiana*

The coding sequences of *PhaTDC1, PhaT6H, PhaT5H, PS1, PS3, PhaPDC1, PDC2-3, BBE1, BBE4, BBE5* were amplified by PCR from cDNA of *P. harmala*, whereas *OMT1, PhaOMT2, OMT3, OMT4,* the *PhaT5H* and *PhaT6H* single and double mutants*, PS2, BBE2_T, BBE2_iso, BBE3, YUC1,* and *CmPDC1* were synthesized by Twist Biosciences. The genes were individually cloned into the pAlpha2-NPT II-Ubq10-CCD-Ter10 vector, which had been digested with ApaI, using Gibson Assembly (Gibson et al., 2009). All constructs were sequenced-verified and subsequently transformed into *Agrobacterium tumefaciens* (*A. tumefaciens*) strain GV3101 by electroporation. Recombinant *A. tumefaciens* carrying overexpression constructs were cultured overnight at 28 °C in LB medium supplemented with kanamycin and gentamicin (50 µg mL^-1^ each). For expression of the mutated *AroG* (AroGF209A, carrying the F209A substitution), previously generated GV3101 stocks kindly provided by Prof. Gad Galili, Prof. Vered Tzin, and Dr. Moran Oliva were used (Tzin et al., 2013). Following overnight incubation, bacterial cells were harvested by centrifugation, washed, and resuspended in infiltration buffer (2 mM Na_3_PO_4_, 50 mM MES, 10 mM MgCl_2_, 0.5 % w v^-1^ glucose, and 100 µM acetosyringone) to an OD_600_ of 1.0. Strains carrying different constructs were combined in equal or defined volumes to achieve the desired gene combinations. The resulting suspensions were infiltrated into the abaxial side of 4-5-week-old *N. benthamiana* leaves using a 1-mL needleless syringe. Leaves were harvested 5-7 days post-infiltration for metabolite analysis. Samples were flash-frozen and extracted as previously described followed by LC-MS/MS analysis.

### Heterologous expression in yeast

The coding sequences of *PhaTDC1*, *PhaT6H*, *PhaPDC1*, and *AtR1* were individually cloned into pESC vector backbones (Agilent Technologies, Santa Clara, CA, USA) to enable dual gene expression under bidirectional, galactose-inducible *GAL1* and *GAL10* promoters. *PhaTDC1* and *PhaT6H* were cloned into pESC-URA under the control of *GAL1* and *GAL10* promoters, respectively, whereas *PhaPDC1* and *AtR1* were cloned into pESC-HIS under the corresponding promoters. Vectors were linearized by PCR, and constructs were assembled *via* In-Fusion cloning. Construct integrity was confirmed by Sanger sequencing before transformation into *S. cerevisiae* strain BY4743 using the Frozen-EZ Yeast Transformation II Kit (Zymo Research, Irvine, CA, USA) according to the manufacturer’s instructions. Yeast co-transformed with empty pESC-HIS and pESC-URA vectors served as the control. Transformants were selected on Synthetic Defined (SD) minimal medium lacking uracil and histidine (SD-URA-HIS), prepared using Yeast Nitrogen Base without amino acids (Sigma-Aldrich, Rehovot, Israel) and appropriate Synthetic Complete drop-out supplements (Formedium, Hunstanton, UK) supplemented with 2% (w v^-1^) D-glucose. Positive colonies were screened and verified by colony PCR. For expression studies, primary cultures were inoculated into 10 mL of SD minimal medium containing 2% (w v^-1^) D-glucose and grown for 24 h at 30 °C with shaking at 200 rpm. To induce heterologous expression, cells were harvested, washed, transferred to SD minimal medium supplemented with 2% (w v^-1^) D-galactose, and incubated for an additional 24 h at 30 °C with shaking at 200 rpm. Cultures were then centrifuged at 4,000 × *g* for 5 min, and cell pellets were collected and weighed. For metabolite extraction, two stainless steel beads (4 mm diameter) and 400 µL of cold methanol were added to the pellets. Samples were homogenized by brief vortexing, sonicated for 20 min, and centrifuged at 15,000 × *g* for 10 min. The resulting supernatant was filtered through a 0.22-µm membrane prior to LC-MS/MS analysis.

### Hairy root transformation of *P. harmala*

The *PvTDC2* and *PvNMT1* genes were amplified from constructs containing the respective genes from Psychotria viridis, as described in our previous study (Berman et al., 2026). Respective binary constructs encoding RFP or RFP::PvNMT1::PvTDC2 were individually assembled into the pAlpha2-NPT II-Ubq10-CCD-Ter10 vector, each under the control of a 35S promoter and NOS terminator, using GoldenBraid cloning (Sarrion-Perdigones et al., 2011). To generate transgenic hairy roots of *P. harmala*, approximately two-week-old cotyledons were infected with *Rhizobium rhizogenes* (*R. rhizogenes*) strain MSU440 harboring the respective binary constructs. Electrocompetent MSU440 cells were transformed by electroporation and cultured in LB medium supplemented with the appropriate antibiotics. Overnight cultures were transferred to LB containing 100 µM Acetosyringone and incubated at 28 °C overnight with shaking (160 rpm). Cells were then pelleted, resuspended in co-cultivation or transformation (CT) medium containing D-glucose (22 g L^-1^) and KH_2_PO_4_ (0.17 g L^-1^, pH 5.5), supplemented with 50 µM Acetosyringone, and adjusted to an OD_600_ of 0.5. Cotyledons were immersed in the *R. rhizogenes* suspension for 30 min, bottled dry, and placed on CT medium containing 100 µM Acetosyringone and 6 g L^-1^ Plant Agar. Explants were co-cultivated with *R. rhizogenes* for 2 days at 23 ± 1°C in the dark. Subsequently, explants were transferred to Murashige and Skoog (MS) medium containing cefotaxime (250 mg L^-1^) and 8 g L^-1^ Plant Agar and incubated at 25 ± 1 °C under partial light. Explants were sub-cultured onto fresh MS medium with appropriate antibiotics every two weeks until transgenic hairy roots emerged, which were then transferred to liquid Gamborg’s B5 medium supplemented with 4% sucrose and the appropriate antibiotics. The B5 medium was refreshed every two weeks.

### Structural modeling and rationale-guided mutagenesis of PhaCYPs

PhaT6H and PhaT5H proteins were modeled using AlphaFold3 (Abramson et al., 2024) with the substrates 6-HT **6** and 5-HT **9**, respectively. AlphaFold3 was installed locally following the developers’ instructions (https://github.com/google-deepmind/alphafold3). Predictions were performed using multiple random seeds, and the plDDT scores of the obtained AlphaFold3 models were in the range of 0.96 to 0.97. Structural models were visualized in PyMOL, and comparison of the predicted active sites identified two divergent residues: Leu373 in PhaT6H versus Val371 in PhaT5H and Thr490 versus Ala488, respectively. These residues were selected for rationale-guided site-directed mutagenesis to assess their contribution to TAM hydroxylation regioselectivity. Mutant genes were synthesized by Twist Biosciences, and individually cloned into the pAlpha2-NPT II-Ubq10-CCD-Ter10 vector, which had been digested with ApaI, using Gibson Assembly (Gibson et al., 2009). The resulting mutants were functionally characterized by agroinfiltration into *N. benthamiana* as described above.

### Statistical analysis

All statistical analyses were conducted using GraphPad Prism (version 8.0.1). Data are presented as median ± standard deviation (SD) based on at least three independent biological replicates. For comparisons between two independent groups, statistical significance was assessed using a two-tailed unpaired Student’s t-test assuming unequal variances. For analyses involving more than two groups, a one-way ANOVA followed by Tukey’s post hoc multiple-comparison test was applied. When ANOVA could not be performed due to zero variance in one or more groups, significance was instead evaluated using a two-tailed unpaired Student’s t-test with unequal variance. The number of biological replicates, statistical tests applied, and significance thresholds are indicated in the corresponding figure legends.

## Data and code availability

All raw next-generation sequencing data will be deposited in the European Nucleotide Archive (ENA), and the sequences of the active genes reported in this manuscript will be deposited at NCBI GenBank. All data and sequences will be made publicly available upon acceptance of the manuscript.

## Supplemental information

Supplemental Figures, Tables and References are provided as Supplemental Information files.

## Author contributions

L.K.H., A.A., and P.B. conceptualized the study. P.B. and J.H. performed the chemical profiling with the aid of I.R. E.A.-S. performed the Iso-Seq transcriptome assembly and annotation, and A.F. performed the TruSeq transcriptome assembly and annotation. L.K.H., J.H., S.C. and P.B. performed all the molecular biology work, *in vitro* enzyme assays, protein purification, and transient expression of genes in *N. benthamiana*. L.K.H. analyzed LC-MS data with the assistance of P.B. L.K.H. and A.K. performed GC-MS analysis of PhaPDC1 activity in *N. benthamiana*. S.C. cloned the genes, performed yeast transformation, and conducted the enzymatic activity assays. Y.D. prepared samples and performed the MALDI imaging experiments. O.K. performed the molecular modeling, analyzed the data, and designed the site-directed mutants. L.K.H. cloned, expressed, and tested the site-directed mutants with the help of S.C. S.C. cloned the genes for the hairy root transformation, B.A.T. performed transformations and generated calli, and L.K.H. maintained the cultures and generated hairy roots. L.K.H. wrote the original draft, and L.K.H. and P.B. reviewed and edited the manuscript with input from all coauthors. S.M., U.H., S.J.F., and I.M. contributed materials and guidance to specific parts of the study. P.B. and A.A. supervised the study and acquired funding. All the authors read and approved the final version of the manuscript.

## Acknowledgments

L.K.H was partially supported by a Faculty Dean fellowship from Faculty of Biochemistry at the Weizmann Institute of Science. A.A. is the incumbent of the Peter J. Cohn Professorial Chair. This study was supported by the Israel Science Foundation (P.B.); and the Adelis Foundation, the Leona M. and Harry B. Helmsley Charitable Trust, the Jeanne and Joseph Nissim Foundation for Life Sciences, Tom and Sondra Rykoff Family Foundation Research, Ron Sklare and the Raymond Burton Plant Genome Research Fund (A.A.). We thank Prof. David Nelson (University of Tennessee, USA) for the systematic naming of the CYPs characterized in this study.

