## Supplemental Figures S1-15 for "Discovery of the β-Carboline Biosynthetic Pathway Enables *In Planta* Reconstruction of the Amazonian Brew Ayahuasca"

### Supplemental Information Table of Contents

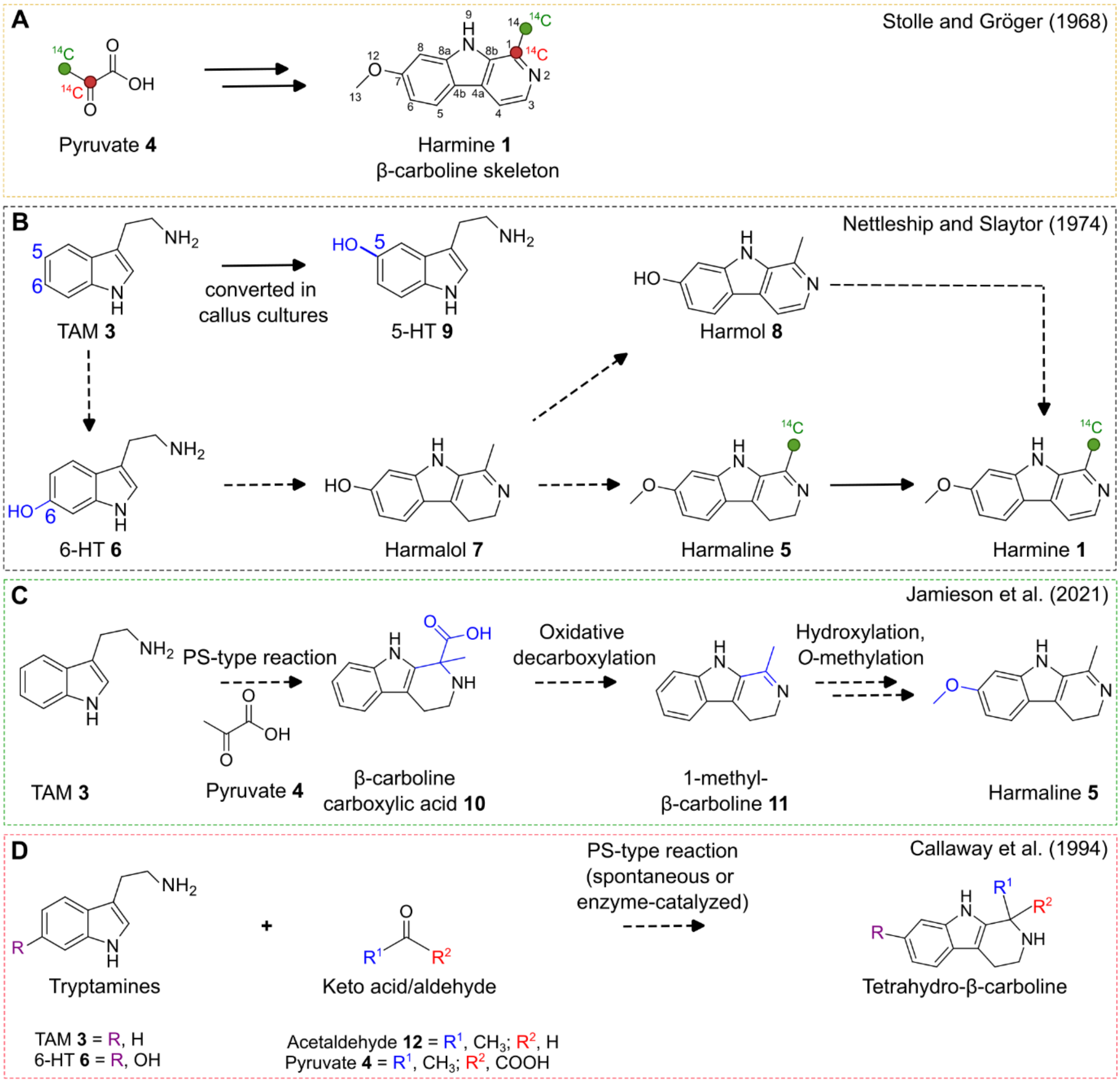

**Supplemental Figure S1. Previously proposed β-carboline biosynthetic pathways in *Peganum harmala* and associated reactions. (A)** Proposed incorporation of pyruvate-derived carbon atoms into the β-carboline skeleton of harmine **1**, based on radiolabeled precursor-feeding experiments (Stolle and Gröger, 1968). Green and red circles indicate the positions of the labeled carbon atoms in pyruvate **4** and their corresponding positions in harmine **1**. **(B)** Previously hypothesized 6-hydroxy-tryptamine (6-HT **6**)-derived pathway for β-carboline biosynthesis. In *Peganum harmala* callus cultures, tryptamine (TAM **3**) was converted to 5-hydroxy-tryptamine (5-HT **9**), whereas feeding with 5-HT **9**, 6-HT **6**, or harmalol **7** did not increase the accumulation of other β-carbolines. In contrast, radio-labeled harmaline **5** was incorporated into harmine **1** (Nettleship and Slaytor, 1974). Green circles indicate the positions of radiolabeled carbon atoms in harmaline **5** and their corresponding positions in harmine **1**. The continuation of **Supplemental Figure S1** appears on the next page.

**(C)** Proposed biosynthetic pathway initiated by Pictet-Spengler (PS)-type condensation of TAM **3** with pyruvate **4**, followed by oxidative decarboxylation, hydroxylation, and *O*-methylation to produce harmaline **5** (Jamieson et al., 2021). **(D)** Potential tetrahydro-β-carboline products formed through PS-type cyclization of tryptamines with pyruvate **4** or acetaldehyde **12** (Callaway et al., 1994). Solid arrows indicate proposed steps supported by experimental evidence in the respective studies, whereas dashed arrows indicate steps that were proposed but not experimentally validated.

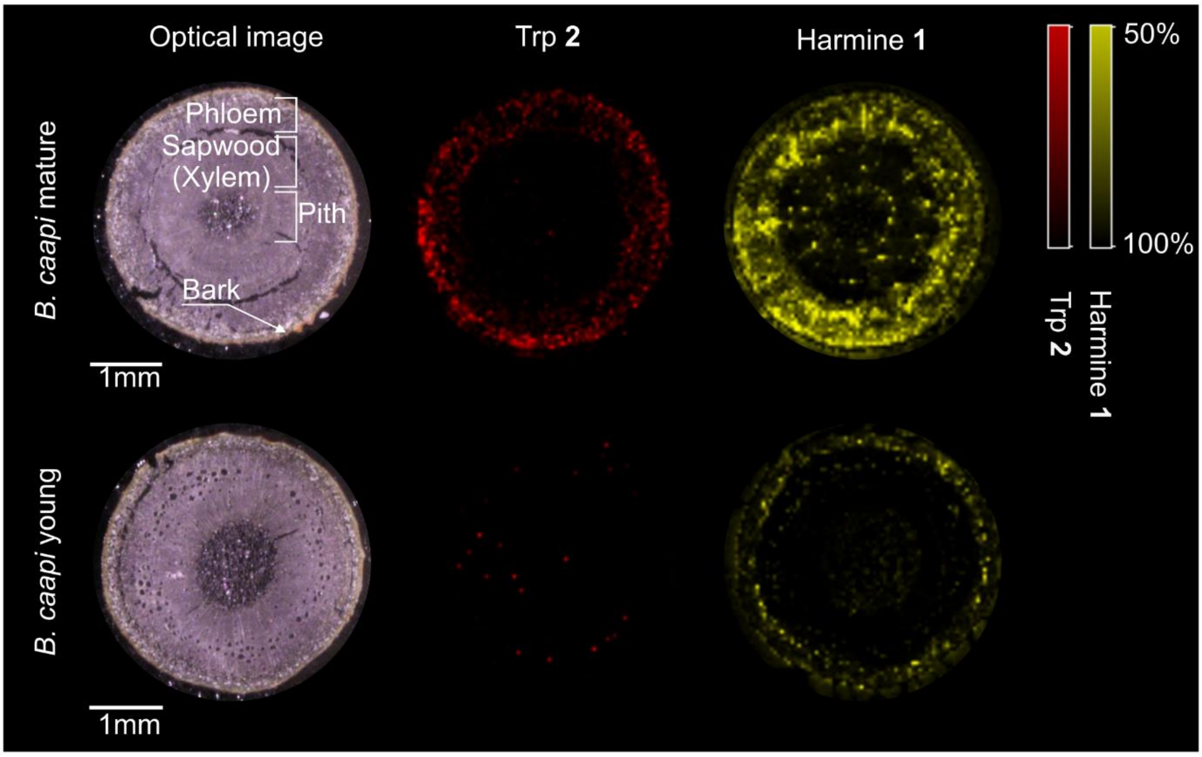

**Supplemental Figure S2. Matrix-assisted laser desorption/ionization–mass spectrometry imaging (MALDI-MSI) of mature and young *Banisteriopsis caapi* stems.** Optical images and MALDI-MSI distributions of tryptophan (Trp **2**, $[M+H]^+$, m/z 205.097 ± 0.01 Da) and harmine **1** ($[M+H]^+$, m/z 213.102 ± 0.01 Da) in stem cross-sections from mature (top) and young (bottom) *Banisteriopsis caapi* tissues ($n$ = 2). Bark, phloem, sapwood (xylem), and pith are indicated in the optical image of the mature stem. Scale bars, 1mm.

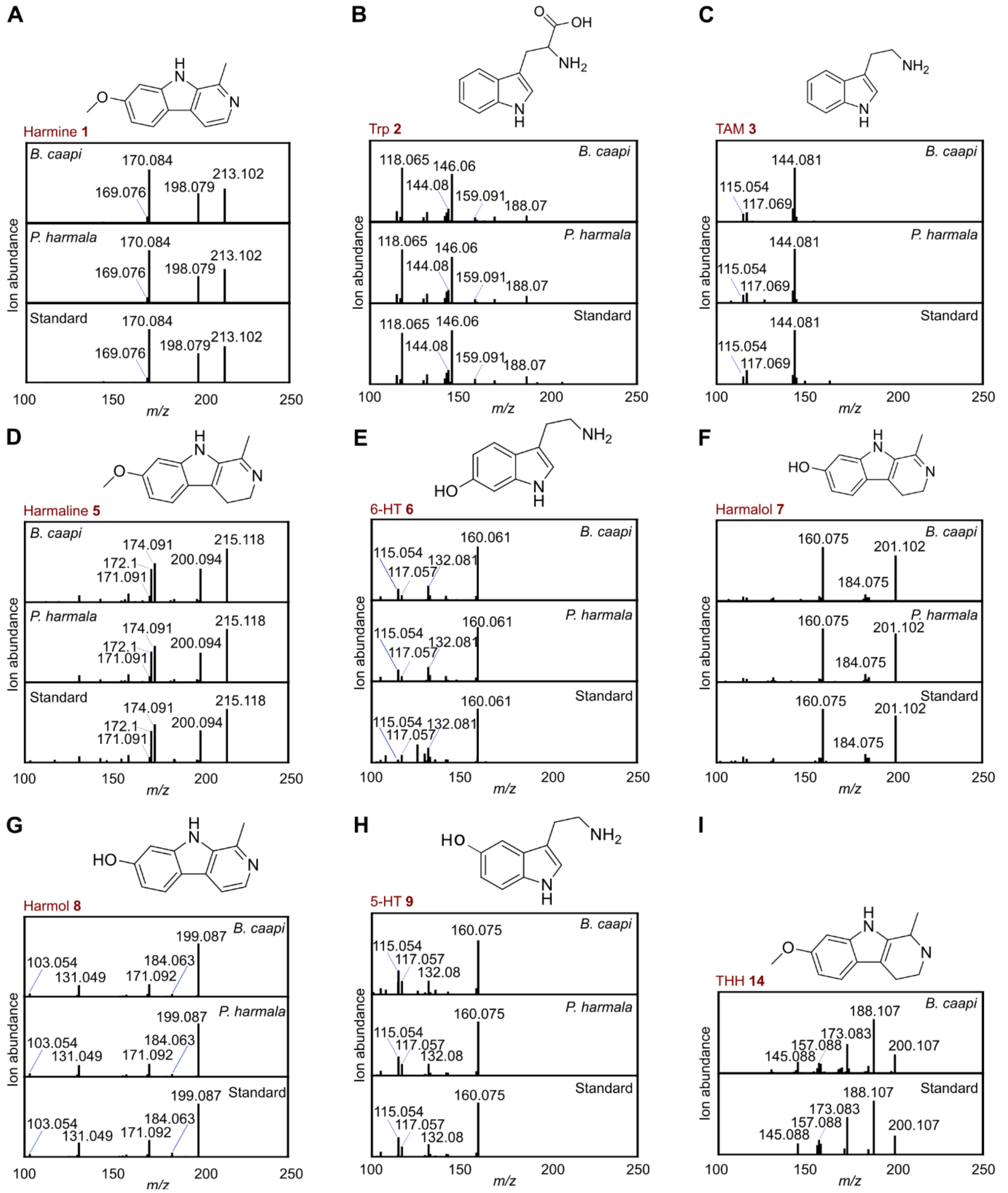

**Supplemental Figure S3. Identification of β-carbolines and intermediates from *Banisteriopsis caapi* and *Peganum harmala* by analytical standards. (A-I)** Tandem mass spectra (MS/MS) spectra matching of metabolites identified in *Banisteriopsis caapi* bark or *Peganum harmala* root extracts versus analytical standards. All the metabolites were identified by LC-MS using analytical standards (full details of each metabolite appear in **Supplemental Table S1)**. Trp **2**, tryptophan; TAM **3**, tryptamine; 6-HT **6**, 6-hydroxy-tryptamine; 5-HT **9**, 5-hydroxy-tryptamine; THH **14**, tetrahydroharmine.

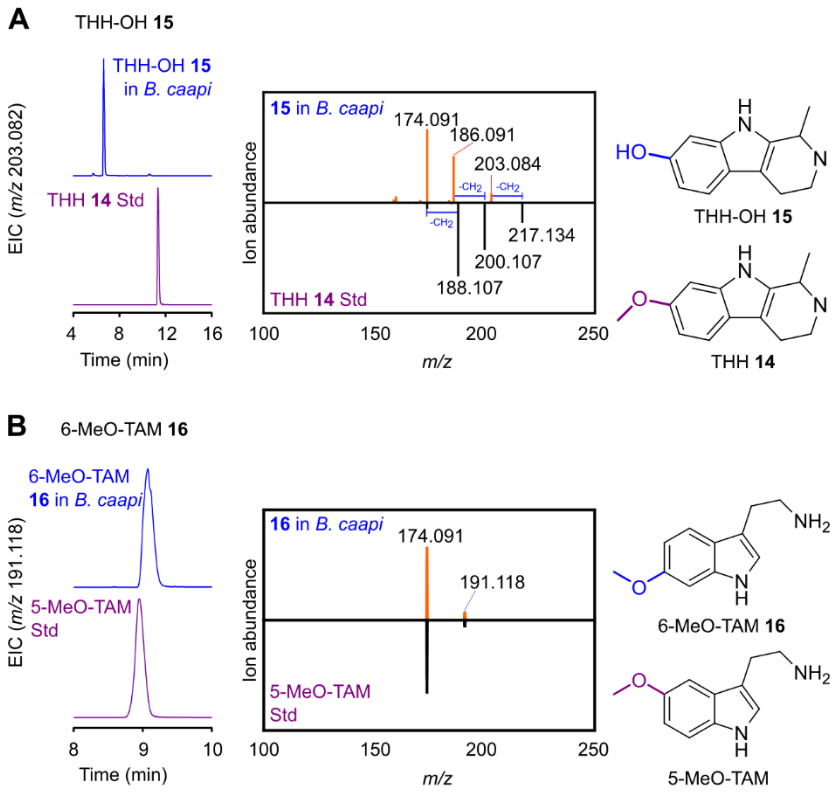

**Supplemental Figure S4. Tentative identification of tetrahydroharmol (THH-OH 15) and 6-methoxy-tryptamine (6-MeO-TAM 16) in *Banisteriopsis caapi*.** Extracted ion chromatograms (EICs) and tandem mass spectra (MS/MS) of metabolites detected in *Banisteriopsis caapi* stem extracts, compared with structurally related compounds for which authentic analytical standards were available. Tentative assignments were based on accurate mass, MS/MS fragmentation patterns, and relative retention times. **(A)** Comparison of THH-OH **15** with tetrahydroharmine (THH **14**). THH-OH **15** exhibited a fragmentation pattern similar to that of THH **14**, with mass differences consistent with the replacement of a methoxy group by a hydroxyl group. Its shorter retention time relative to THH **14** was consistent with its greater predicted polarity, further supporting the tentative assignment. **(B)** Comparison of 6-MeO-TAM **16** with an authentic 5-methoxytryptamine (5-MeO-TAM) standard. The tentative identification of 6-MeO-TAM **16** was supported by its accurate mass and MS/MS fragmentation pattern, consistent with those of the structurally related 5-MeO-TAM standard, and by its chromatographic behavior, with the relative elution order of the two regioisomers matching that observed for 5-hydroxy-tryptamine (5-HT **9**) and 6-hydroxy-tryptamine (6-HT **6**; retention times for all the metabolites appear in **Supplementary Table S1**). Corresponding chemical structures are included for reference, with the relevant functional groups highlighted.

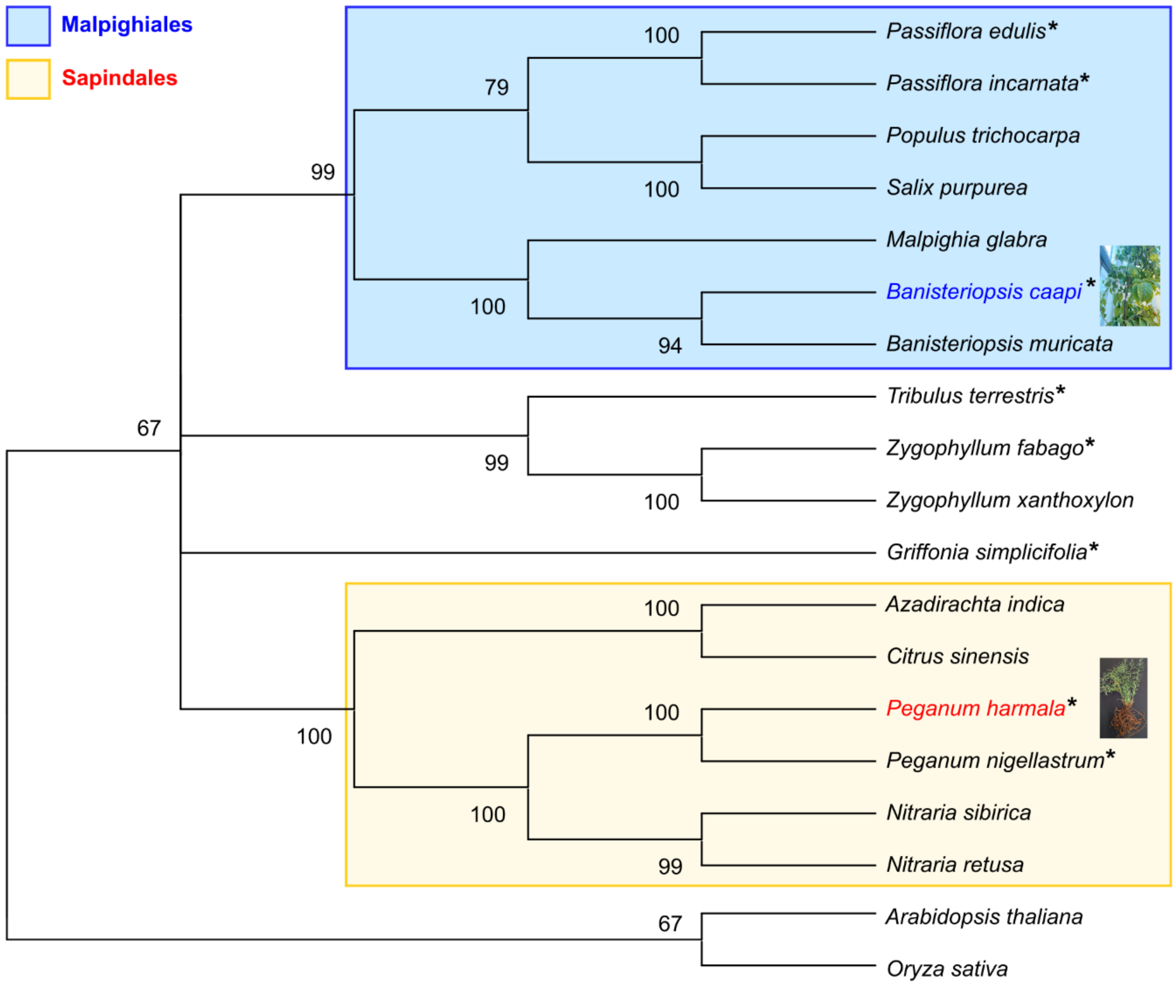

**Supplemental Figure S5. Phylogenetic relationships of β-carboline-producing plants.** The tree was constructed using amino acid sequences of maturase K (MatK), encoded by the chloroplast *matK* gene. Representative plant species are included, with β-carboline-producing plants marked with asterisks (Fritzsche, 1848; Slaytor and McFarlane, 1968; Schultes, 1975; Tsuchiya et al., 1999; Grice et al., 2001; Wang et al., 2009; Wang et al., 2013; López-Orenes et al., 2014). The plant orders Malpighiales and Sapinadales which include major β-carboline-producing lineages represented in this analysis, are highlighted in blue and yellow, respectively. *Arabidopsis thaliana* and *Oryza sativa* were included as reference species. A full list of GenBank accession numbers appears in **Supplemental Table S3**. The rooted maximum-likelihood tree was constructed using a MUSCLE multiple alignment in MEGA12 software, with 1,000 bootstrap replicates; bootstrap values are indicated at the nodes.

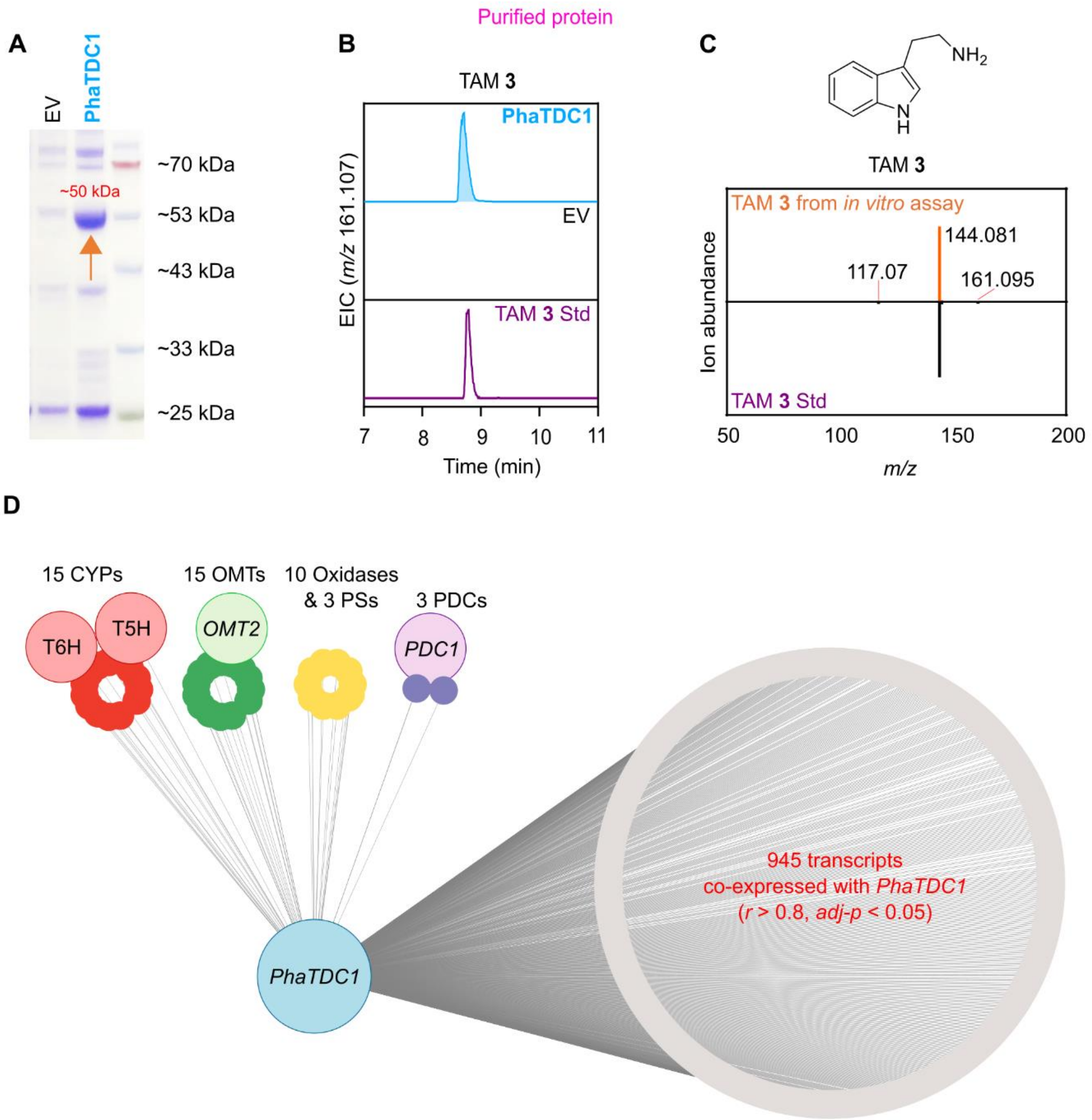

**Supplemental Figure S6. *In vitro* characterization of PhaTDC1 and co-expression analysis.** *In vitro* functional characterization of PhaTDC1, showing (**A**) gel electrophoresis of the purified PhaTDC1 with the corresponding protein band indicated by an arrow, **(B)** extracted ion chromatograms of tryptamine (TAM **3**) produced in assays with purified *Escherichia coli*-expressed *PhaTDC1* or an empty-vector control in the presence of tryptophan (Trp **2**) and pyridoxal 5′-phosphate (PLP) ($n$ = 3 biological replicates), and (**C**) tandem mass spectra (MS/MS) of the produced TAM **3** compared with an authentic analytical standard. Chromatograms were normalized to the highest values. **(D)** Co-expression network of iso-seq transcripts correlated with *PhaTDC1* (visualized using Cytoscape). Pairwise Pearson correlation coefficients were calculated between *PhaTDC1* and transcripts across analyzed samples. The continuation of **Supplemental Figure S6** appears on the next page.

Transcripts with a threshold of Pearson correlation coefficient (*r*) > 0.8 and false discovery rate (FDR)-adjusted *p*-values < 0.05 were considered co-expressed. The network includes 15 cytochrome P450s (CYPs), 3 pyruvate decarboxylases (PDCs), 3 Pictet-Spenglerases (PSs), 15 *O*-methyltransferases (OMTs), and 10 oxidases. The total number of co-expressed transcripts and functionally characterized active enzymes identified in this study are indicated. Edge width represents the absolute value of the Pearson correlation coefficient. EV, empty vector; Std, standard.

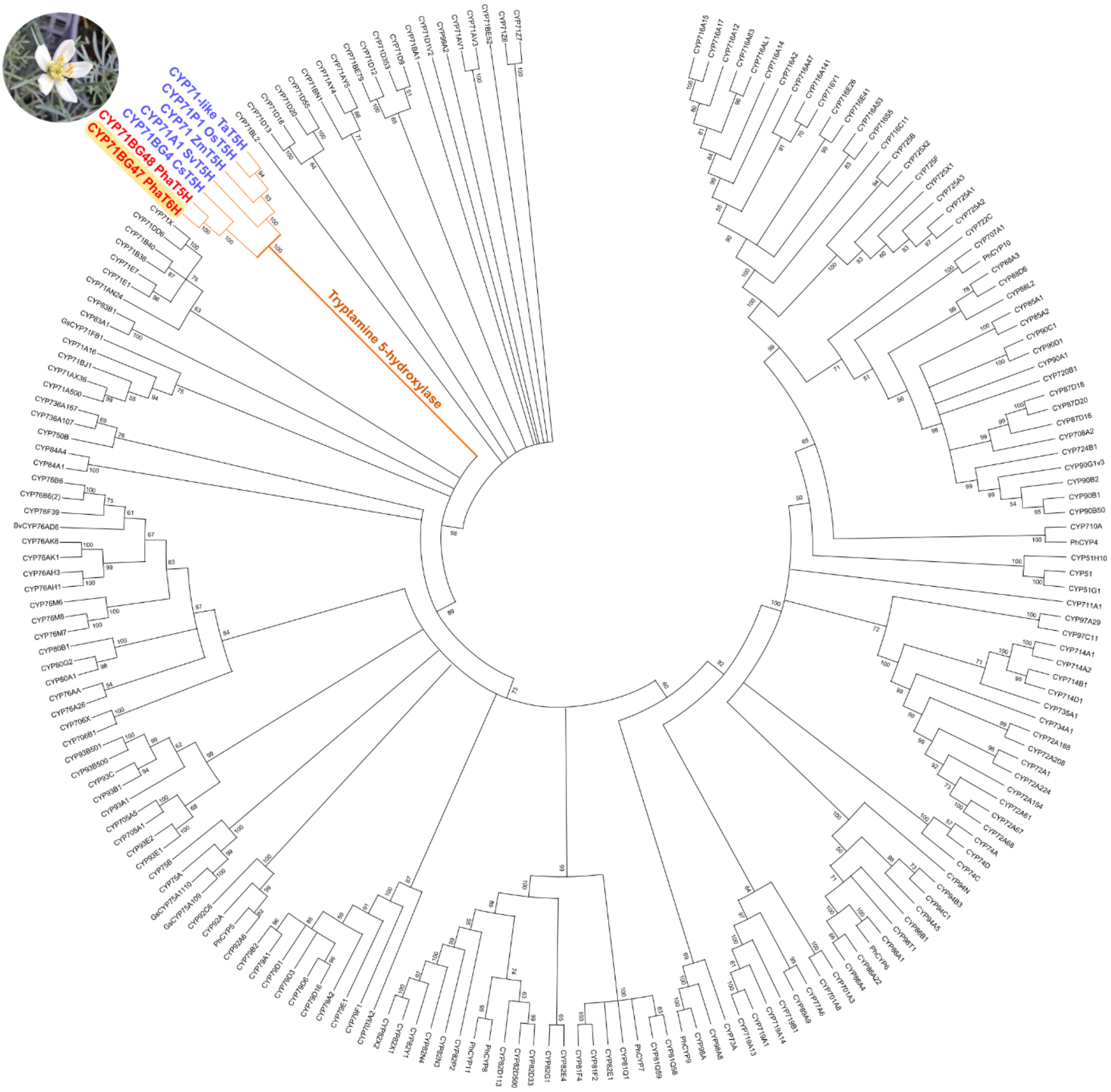

**Supplemental Figure S7. Phylogenetic analysis of *Peganum harmala* cytochrome P450 enzymes.** Putative CYP proteins from *P. harmala* (marked in red) and functionally characterized plant CYPs (Plant cytochrome P450 database, 2020) are shown. Previously characterized tryptamine-5-hydroxylase (T5H) enzymes are marked in blue. Cs, *Citrus sinensis*; Sv, *Setaria viridis*; Zm, *Zea mays*; Os, *Oryza sativa*; Ta, *Triticum aestivum*. The full list of protein IDs is provided in **Supplemental Table S6**. The maximum likelihood tree was constructed with 1000 bootstrap tests on the basis of a MUSCLE multiple alignment using MEGA12 software; bootstrap values are indicated at the nodes.

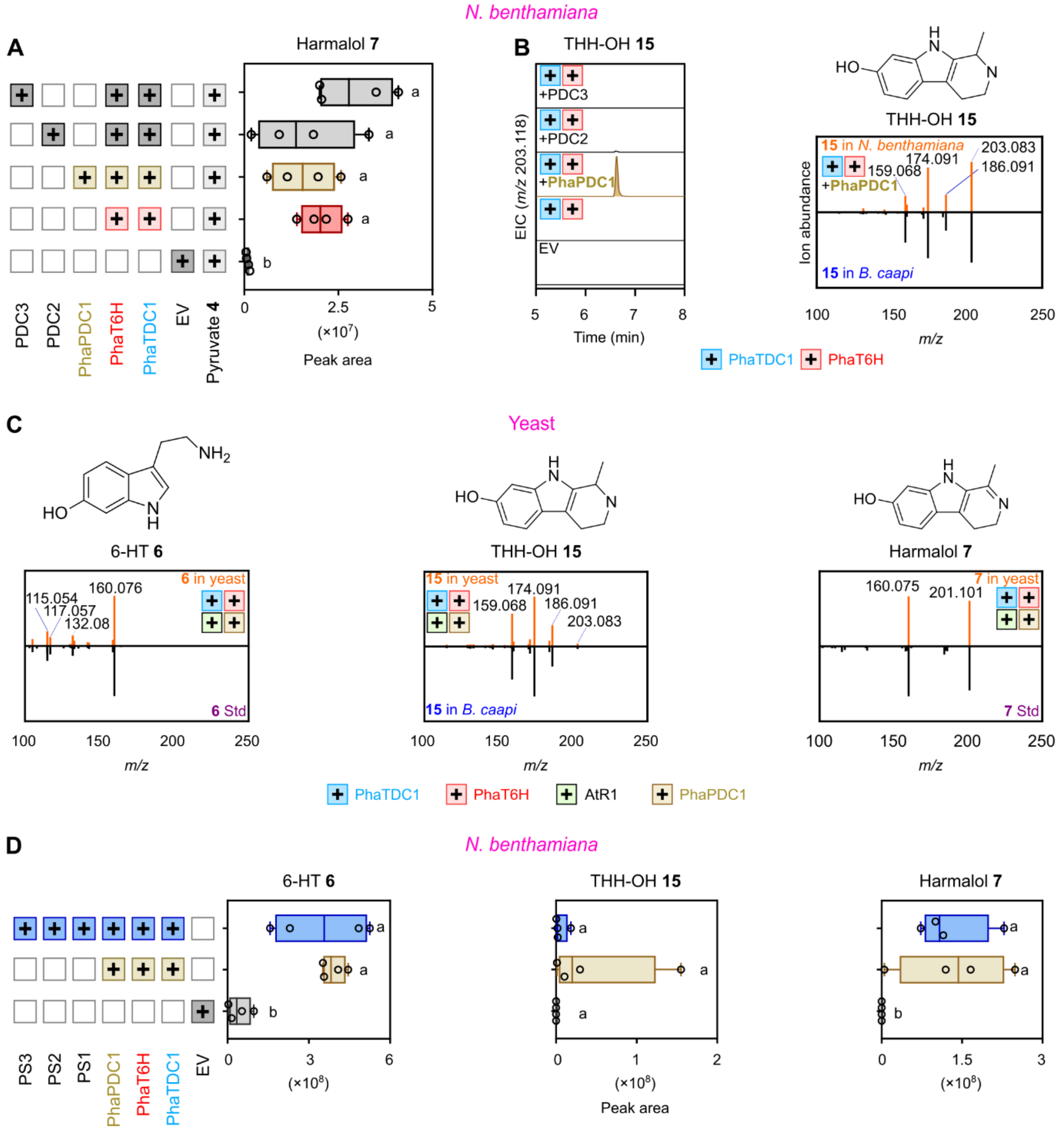

**Supplemental Figure S8. *In planta* assays of pyruvate decarboxylase (PDC) and pictet-spenglerase (PS) candidates in *Nicotiana benthamiana*. (A)** Ion abundance peak areas (median ± SD; $n$ = 4 biological replicates), and **(B)** extracted ion chromatograms (EICs) and tandem mass spectra (MS/MS) of harmalol **7** and tetrahydroharmol (THH-OH **15**), respectively, following transient co-expression of *PhaTDC1*, *PhaT6H*, *PhaPDC1,* and *PDC2-3* candidates in *Nicotiana benthamiana* (*N. benthamiana*) leaves. The EIC of THH-OH **15** from *Banisteriopsis caapi* (*B. caapi*) stem extract is shown as reference in the absence of an analytical standard. Chromatograms were normalized to the highest values. The continuation of **Supplemental Figure S8** appears on the next page.

**(C)** MS/MS spectra matching of 6-hydroxy-tryptamine (6-HT **6**), THH-OH **15**, and harmalol **7** following heterologous co-expression of *PhaTDC1*, *PhaT6H*, with or without *PhaPDC1*, together with *AtR1* in *Saccharomyces cerevisiae*. **(D)** Ion abundance peak areas (median ± SD; $n$ = 4 biological replicates) of 6-HT **6**, THH-OH **15**, and harmalol **7** following transient expression of *PhaTDC1*, *PhaT6H*, *PhaPDC1,* and *PS1-3* candidates in *N. benthamiana* leaves. Statistical significance was assessed by one-way ANOVA followed by Tukey's HSD test, letters denote statistically significant differences ($p < 0.05$). Plus signs (+) in **(A)** and **(D)** indicate biosynthetic genes expressed in each experiment, with boxes colored according to the last enzyme added in a set of coexpressed genes. Metabolites were identified by comparison of their MS/MS spectra with those of analytical standards or reference extracts from *B. caapi*. Leaves from *N. benthamiana* samples were collected 5 days after infiltration. EV, empty vector; Std, standard.

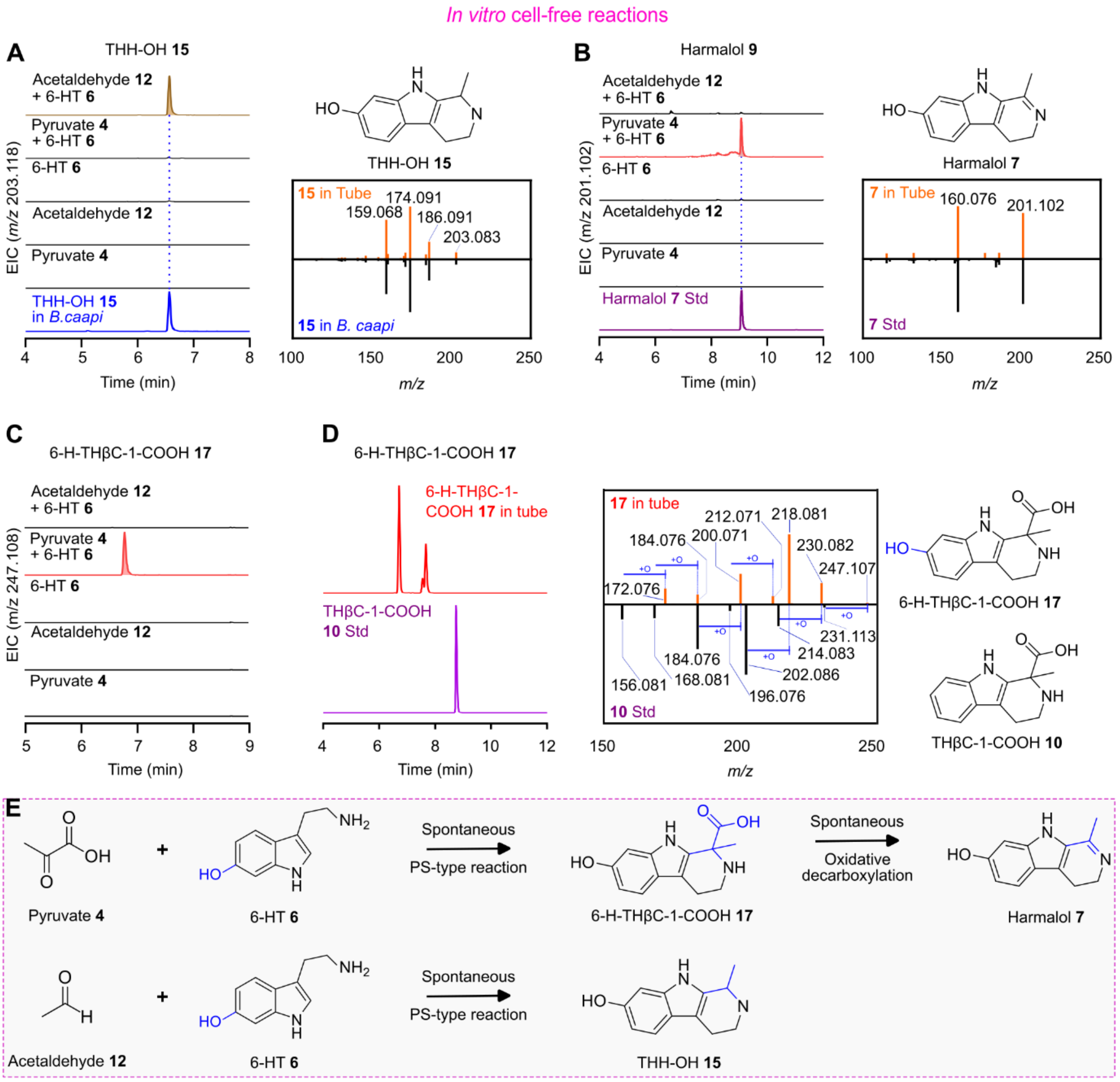

**Supplemental Figure S9. Spontaneous β-carboline scaffold formation in cell-free reactions.** Extracted ion chromatograms (EICs) and tandem mass spectra (MS/MS) of **(A)** tetrahydroharmol (THH-OH **15**), **(B)** harmalol **7**, and **(C)** 6-hydroxy-1,2,3,4-tetrahydro-β-carboline-1-carboxylic acid (6-H-THβC-1-COOH **17**) detected in cell-free reactions containing different combinations of pyruvate **4**, acetaldehyde **12**, and 6-hydroxy-tryptamine (6-HT **6**). Reactions were incubated at 37 °C for 1 h (*n* = 3 biological replicates). Chromatograms were normalized to the highest values. **(D)** EIC and MS/MS spectra of 6-H-THβC-1-COOH **17** detected in cell-free reactions containing pyruvate **4** and 6-HT **6**, compared with those of an authentic 1,2,3,4-tetrahydro-β-carboline-1-carboxylic acid (THβC-1-COOH **10**) analytical standard. The continuation of **Supplemental Figure S9** appears on the next page.

In the absence of an authentic standard, 6-H-THβC-1-COOH **17** was tentatively identified based on its accurate mass and MS/MS fragmentation pattern, which was similar to that of THβC-1-COOH **10**, with a mass shift corresponding with the addition of an oxygen atom through hydroxylation. The shorter retention time of 6-H-THβC-1-COOH **17** relative to THβC-1-COOH **10** was consistent with its greater polarity due to the additional hydroxyl group, further supporting the tentative assignment. Corresponding chemical structures are included for reference. **(E)** Schematic representation of spontaneous β-carboline scaffold formation in cell-free reactions. Pyruvate **4** reacts with 6-HT **6** through a spontaneous Pictet-Spengler (PS)-type reaction to form 6-H-THβC-1-COOH **17**, followed by oxidative decarboxylation to produce harmalol **7**. Alternatively, acetaldehyde **12** undergoes spontaneous PS-type reaction with 6-HT **6** to form THH-OH **15**. RT, mature roots; RY, young roots; Std, standard.

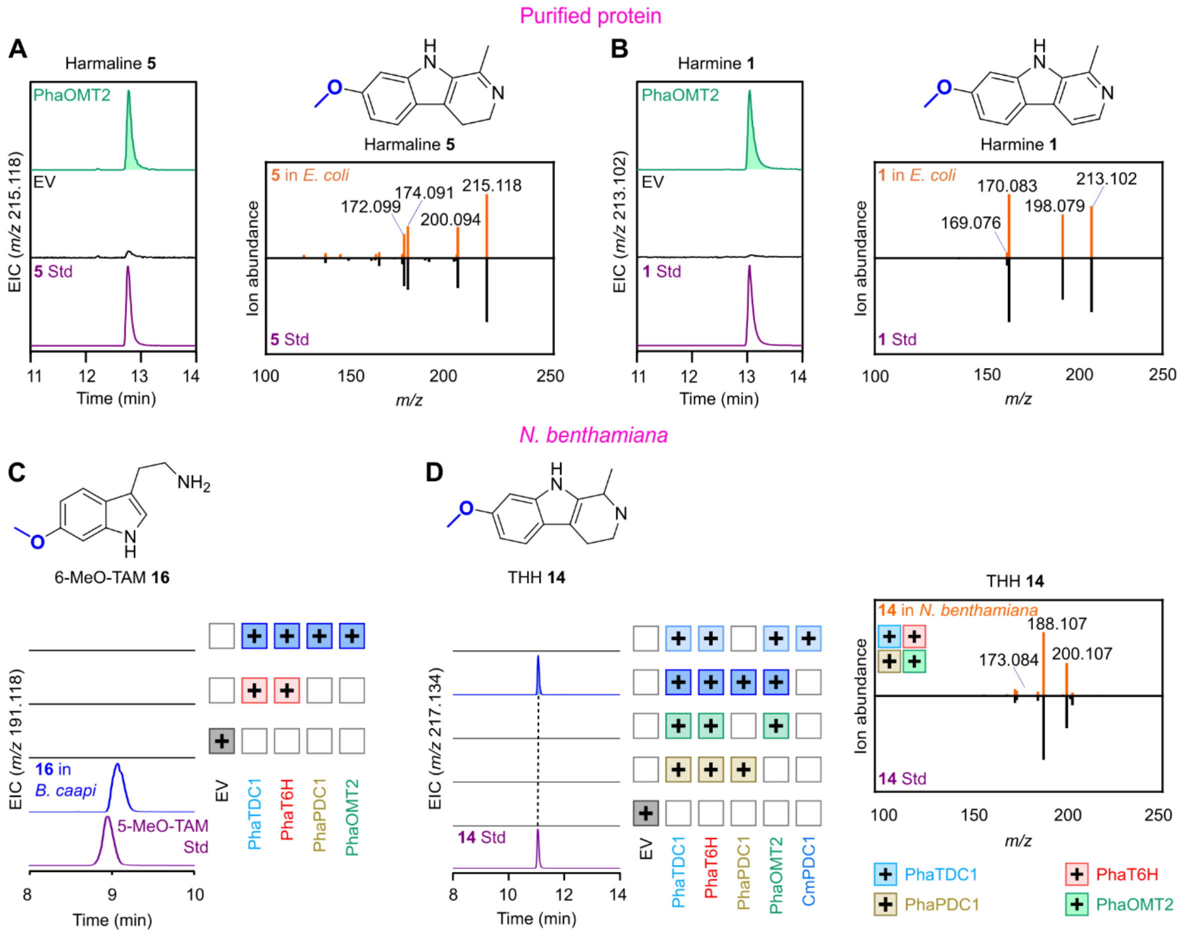

**Supplemental Figure S10. Functional characterization of PhaOMT2. (A-B)** Extracted ion chromatograms (EICs) and tandem mass spectra (MS/MS) of harmaline **5** and harmine **1** produced in *in vitro* enzyme assays with purified *Escherichia coli*-expressed PhaOMT2, with S-adenosyl-L-methionine (SAM) and harmalol **7** or harmol **8** as substrates, respectively (*n* = 3 biological replicates). **(C)** EICs of 6-methoxytryptamine (6-MeO-TAM **16**) detected in *Banisteriopsis caapi* (*B. caapi*) and 5-methoxytryptamine (5-MeO-TAM) analytical standard. No detectable 6-MeO-TAM **16** was observed in *Nicotiana benthamiana* (*N. benthamiana*) leaves following transient expression of *PhaTDC1* and *PhaT6H*, either alone or in combination with *PhaPDC1* and *PhaOMT2*. **(D)** EICs and MS/MS spectra of THH **14** following transient co-expression of *PhaTDC1*, *PhaT6H*, and different combinations of *PhaPDC1* and *PhaOMT2* in *N. benthamiana* leaves. Leaves were collected 7 days after infiltration (*n* = 4 biological replicates). Chromatograms were normalized to the highest values. Spectra were compared to analytical standards. Corresponding chemical structures are included for reference. Plus signs (+) in **(C)** and **(D)** indicate biosynthetic genes expressed in each experiment, with boxes colored according to the last enzyme added in a set of coexpressed genes. THH **14**, tetrahydroharmine; EV, empty vector; Std, standard.

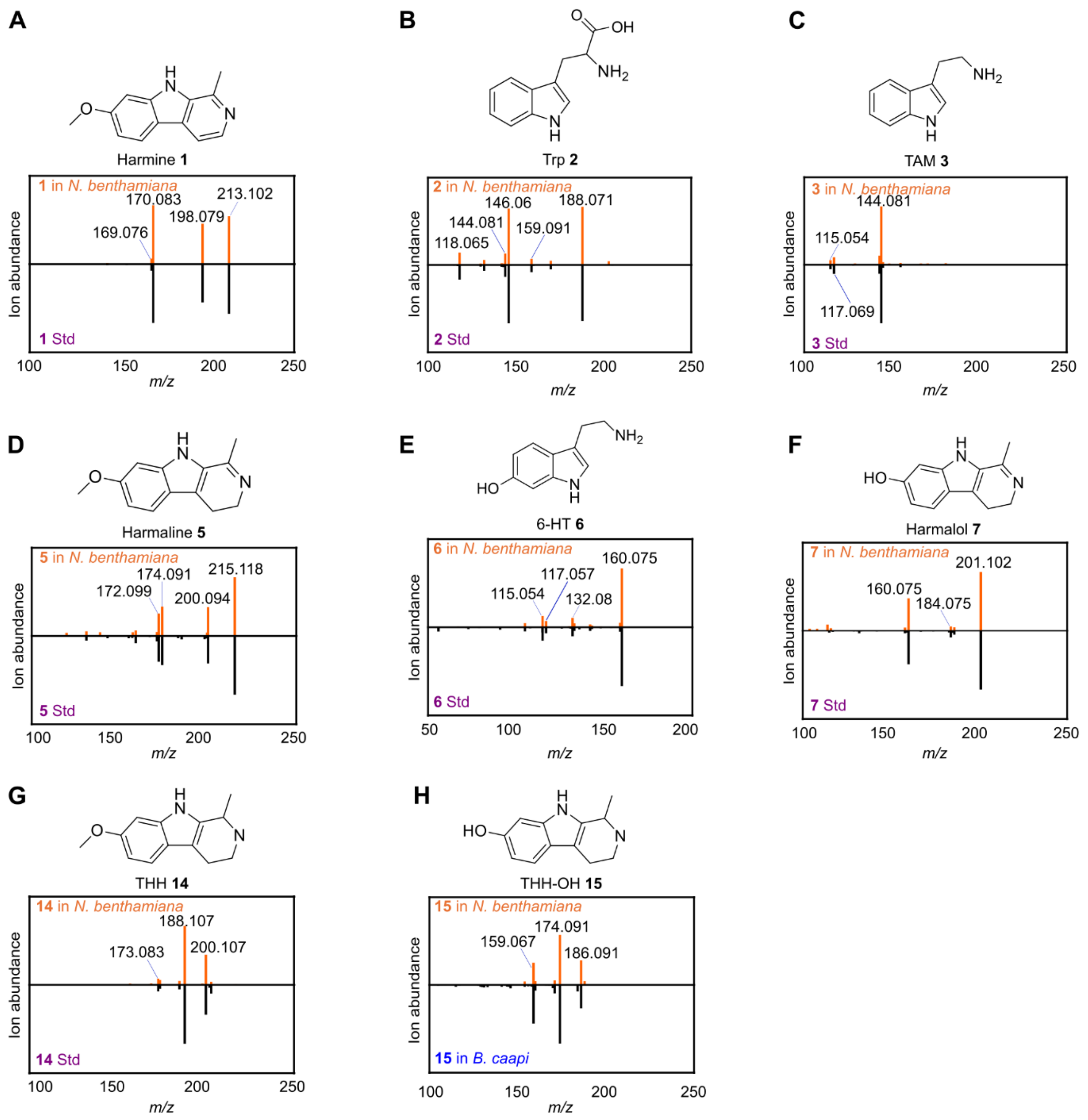

**Supplemental Figure S11. Identification of β-carbolines and biosynthetic intermediates following pathway reconstruction in *Nicotiana benthamiana*. (A-H)** Comparison of tandem mass spectra (MS/MS) of standards and corresponding metabolites detected in *Nicotiana benthamiana* leaf extracts following co-expression of *AroG, PhaTDC1*, *PhaT6H*, *PhaPDC1*, and *PhaOMT2*. Metabolites were identified by comparing retention times, accurate masses, and MS/MS spectra with those of analytical standards or a reference stem extract from *Banisteriopsis caapi* (full details of each metabolite appear in **Supplemental Table S1**). Trp **2**, tryptophan; TAM **3**, tryptamine; 5-HT **9**, 5-hydroxy-tryptamine; 6-HT **6**, 6-hydroxy-tryptamine; THH-OH **15**, tetrahydroharmol; THH **14**, tetrahydroharmine; Std, standard.

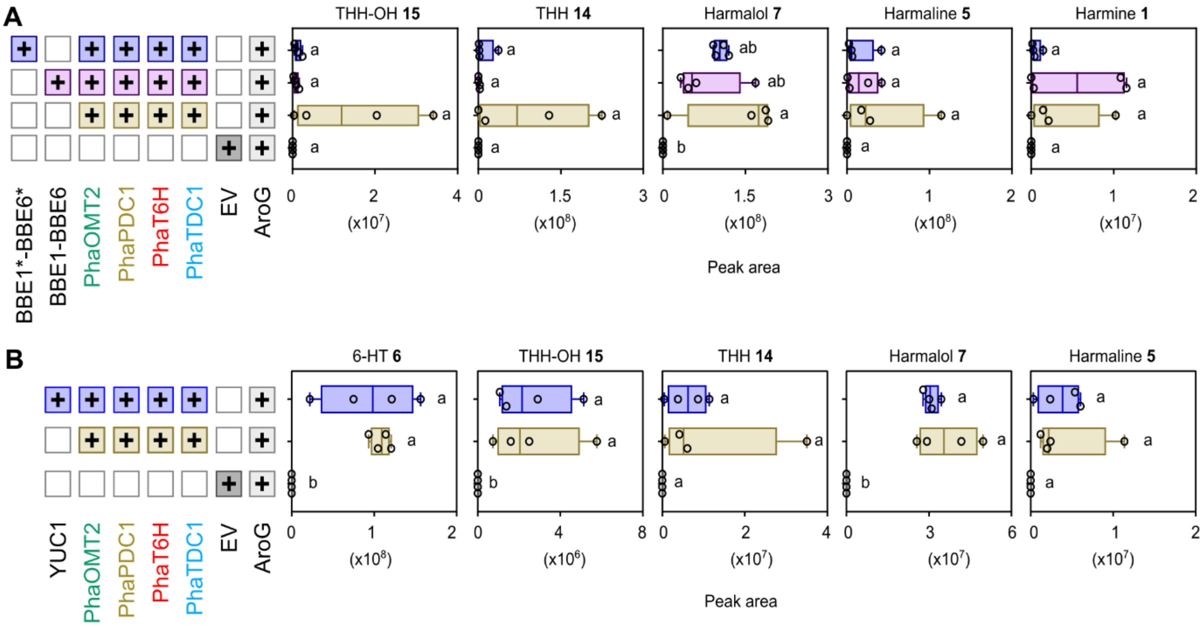

**Supplemental Figure S12. Screening of candidate oxidases potentially involved in β-carboline biosynthesis in *Peganum harmala*.** Ion abundance peak areas (median ± SD; $n = 4$ biological replicates) of β-carbolines following transient co-expression of *AroG*, *PhaTDC1*, *PhaT6H*, *PhaPDC1*, *PhaOMT2* in *Nicotiana benthamiana* leaves, together with **(A)** six *BBE-like* oxidase candidates, expressed with or without their predicted signal peptides (signal peptide deletion marked by an asterisk, *), or **(B)** the *YUC1* candidate. Leaves were collected 7 days after infiltration. Plus signs (+) indicate biosynthetic genes expressed in each experiment, with boxes colored according to the last enzyme added in a set of coexpressed genes. Statistical significance was determined by one-way ANOVA followed by Tukey's HSD post hoc test. Different letters indicate statistically significant differences between groups ($p < 0.05$). 6-HT **6**, 6-hydroxy-tryptamine; THH-OH **15**, tetrahydroharmol; THH **14**, tetrahydroharmine; EV, empty vector.

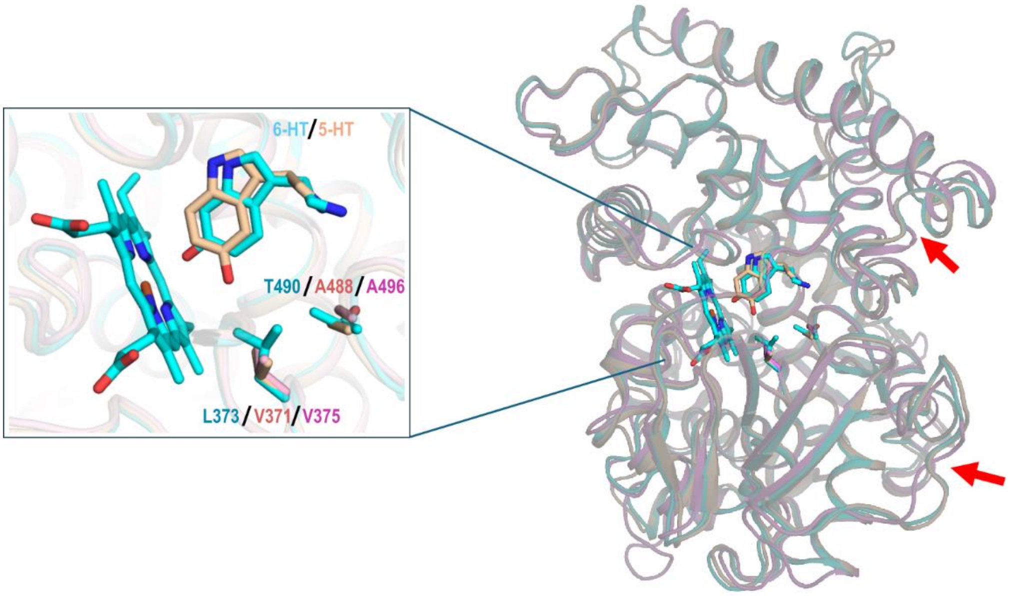

**Supplemental Figure S13. Structural comparison of tryptamine hydroxylases from *Peganum harmala* and *Oryza sativa*.** Structural overlay of 6-hydroxy-tryptamine (6-HT **6**) bound to PhaT6H (cyan), and 5-hydroxy-tryptamine (5-HT **9**) bound to PhaT5H (wheat) and to OsT5H (pink). Heme, ligands, and the two residues differing between the PhaT6H and PhaT5H active sites are shown as sticks, while key backbone differences between the PhaCYPs and OsT5H are indicated by red arrows. L373 and T490 in PhaT6H (corresponding to V371 and A488 in PhaT5H) appear incompatible with 5-HT **9** production.

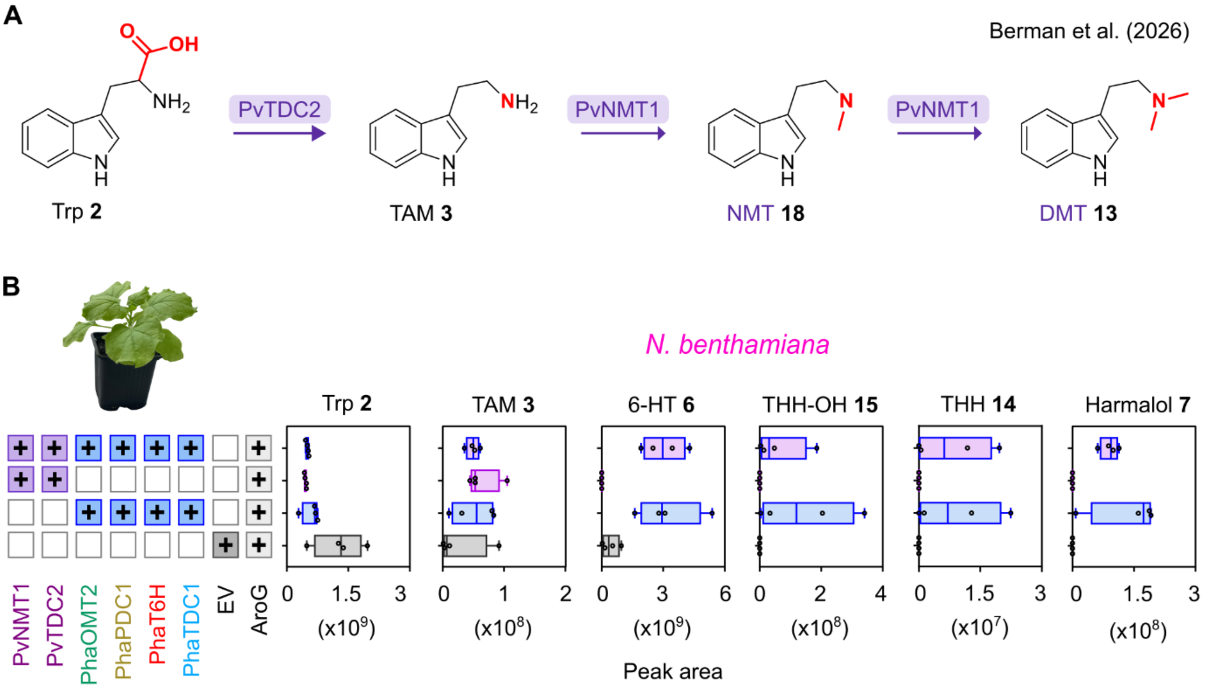

**Supplemental Figure S14. Combined production of β-carbolines and *N,N*-dimethyltryptamine (DMT 13) in *Nicotiana benthamiana*.** (A) Complete biosynthetic pathway of DMT in *Psychotria viridis* (*P. viridis*) (Berman et al., 2026). (B) Representative image of *Nicotiana benthamiana* (*N. benthamiana*) and ion abundance peak areas (median ± SD; n = 4 biological replicates) of β-carboline intermediates detected in *N. benthamiana* leaves following transient expression of harmine **1** biosynthetic genes (*PhaTDC1*, *PhaT6H*, *PhaPDC1*, and *PhaOMT2*), DMT **13** biosynthetic genes (*PvTDC2* and *PvNMT1*) from *P. viridis* (Berman et al., 2026), or their combination. The bacterial feedback-insensitive *AroG* mutant was co-expressed in all combinations to enhance the accumulation of aromatic amino acids in *N. benthamiana* leaves. Samples were harvested 7 days after infiltration. Plus signs (+) indicate biosynthetic genes expressed in each experiment, with blue, purple, and combined blue-purple boxes representing β-carboline, DMT 13, and combined biosynthetic genes, respectively. Trp **2**, tryptophan; TAM **3**, tryptamine; 6-hydroxy-tryptamine, 6-HT **6**; THH-OH **15**, tetrahydroharmol; THH **14**, tetrahydroharmine; Std, standard; EV, empty vector.

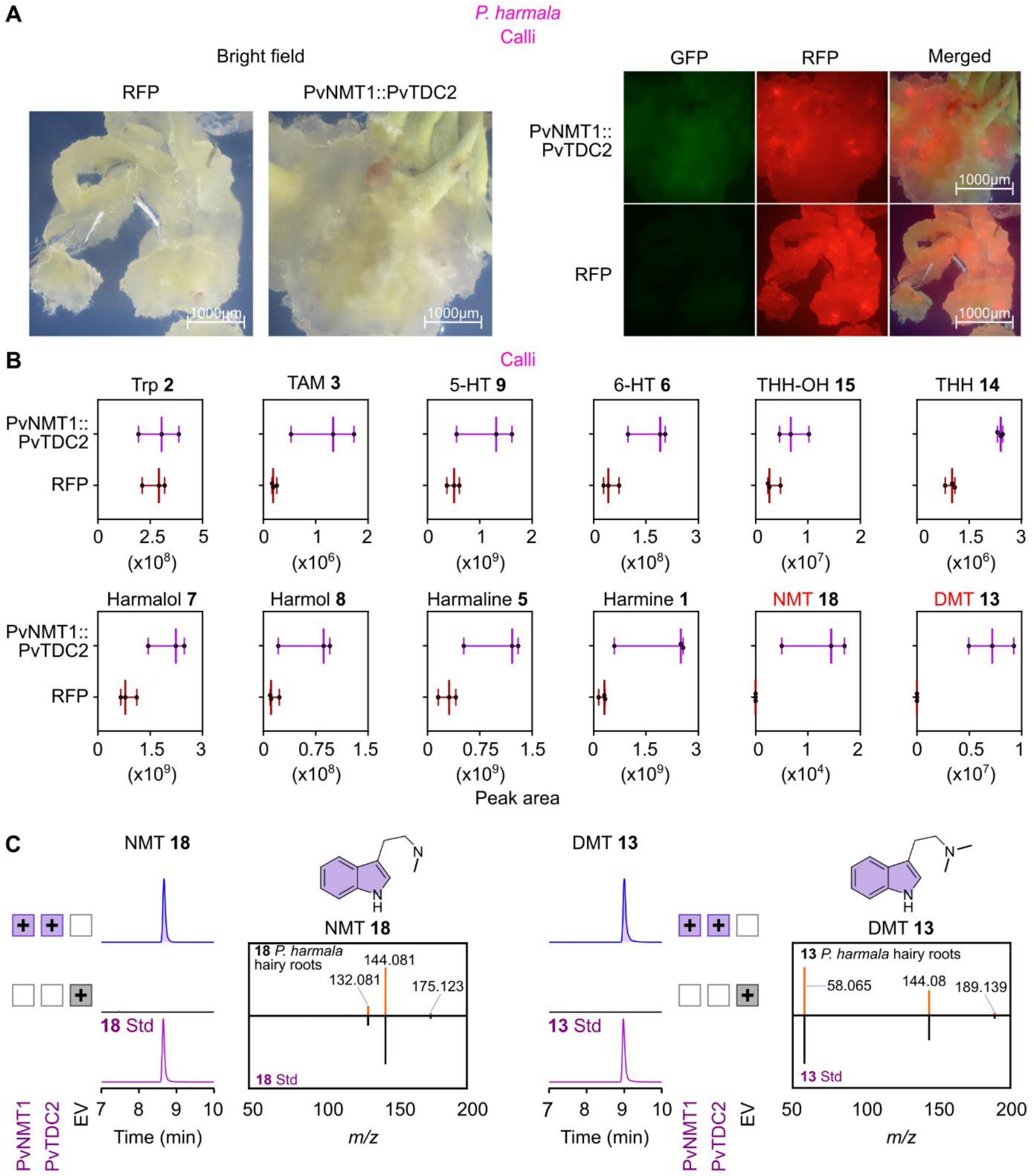

**Supplemental Figure S15. Combined production of β-carbolines and *N,N*-dimethyltryptamine (DMT 13) in a single plant tissue. (A-C)** *Peganum harmala* (*P. harmala)* cotyledons were transformed using *Rhizobium rhizogenes* (MSU440)-mediated transformation with a construct expressing DMT **13** (*PvTDC2* and *PvNMT1*) from *Psychotria viridis* (*P. viridis*) (Berman et al., 2026). Hairy roots or calli transformed with an RFP-expressing vector served as controls. **(A)** Representative bright-field, green fluorescence (GFP) or red fluorescence (RFP), and merged images of transformed *P. harmala* calli. The continuation of **Supplemental Figure S15** appears on the next page.

**(B)** Ion abundance peak areas (median ± SD; $n$ = 3 biological replicates) of harmine **1**, DMT **13**, and their intermediates detected in transgenic calli expressing *PvTDC2*, and *PvNMT1*. **(C)** Extracted ion chromatograms and tandem mass spectra of *N*-methyltryptamine (NMT **18**) and DMT **13** detected in transgenic hairy roots expressing *PvTDC2* and *PvNMT1.* Chemical structures are included for reference. Both compounds were identified by comparing their accurate masses, retention times and MS/MS spectra with those of authentic analytical standards. Trp **2**, tryptophan; TAM **3**, tryptamine; 5-HT **9**, 5-hydroxy-tryptamine; 6-HT **6**, 6-hydroxy-tryptamine; THH-OH **15**, tetrahydroharmol; THH **14**, tetrahydroharmine; Std, standard; EV, empty vector.
